# Hit-and-run silencing of endogenous *DUX4* by targeting DNA hypomethylation on D4Z4 repeat array in facioscapulohumeral muscular dystrophy

**DOI:** 10.1101/2022.04.12.487997

**Authors:** Junjie He, Mitsuru Sasaki-Honda, Tatsuya Jonouchi, Meni Arai, Takumi Kishimoto, Aki Matsumoto, Chihiro Mori, Sayaka Ido, Satoko Sakurai, Takuya Yamamoto, Hidetoshi Sakurai

**Author notes:** Correspondence may also be addressed to Mitsuru Sasaki-Honda. These authors contributed equally to this work (Junjie He, Mitsuru Sasaki-Honda).

## Abstract

Facioscapulohumeral muscular dystrophy (FSHD), a progressive skeletal muscle disorder, is characterized by DNA hypomethylation of the D4Z4 repeat array in the 4q35 region, resulting in aberrant *DUX4* expression. Restoring sustainable *DUX4* repression therefore represents a promising therapeutic strategy to halt disease progression. However, most approaches rely on sustained expression of biochemical entities, raising safety concerns and potentially narrowing the quality of life of FSHD individuals. Here, we demonstrate that transient “hit-and-run” dCas9-mediated epigenetic silencing delivered via RNAs can target D4Z4 hypomethylation. This approach achieved sustained repression of endogenous *DUX4* and reversed disease-associated transcriptional dysregulation in a disease model using patient-derived induced pluripotent stem cells (iPSCs). Notably, in both repeat contraction-dependent FSHD1 and contraction-independent FSHD2, D4Z4 DNA methylation increased robustly, leading to gene repression that persisted for at least two weeks post-transfection and restoration of disease-associated pathways, while transient delivery of conventional CRISPR interference didn’t show comparable durability. Notably, transient RNA delivery of the editing tools minimized the global off-target impact on the transcriptome and methylome associated with constitutive expression of epigenetic editors. These findings establish hit-and-run silencing targeting congenital DNA hypomethylation for sustained gene suppression, supporting its therapeutic potential for diseases driven by pathogenic DNA hypomethylation or repeat-associated epigenetic dysregulation.

**GRAPHICAL ABSTRACT:** 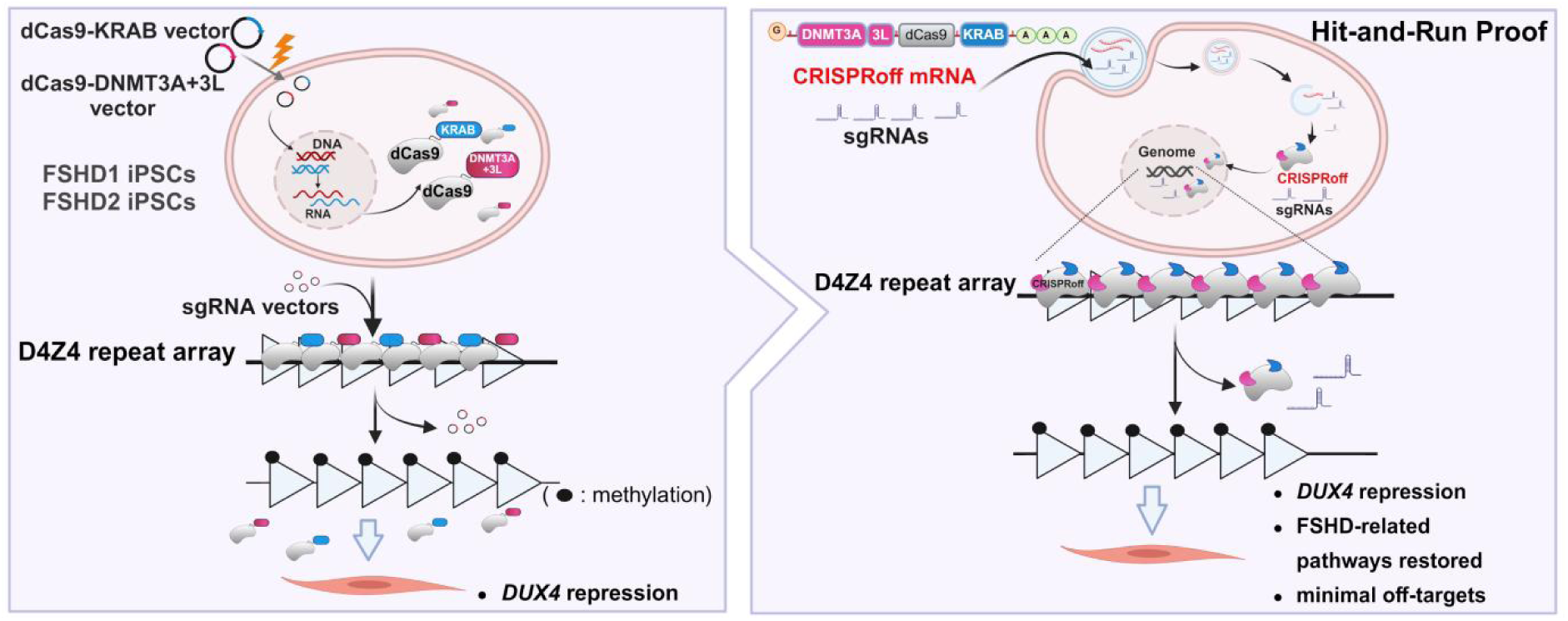

## INTRODUCTION

Facioscapulohumeral muscular dystrophy (FSHD, OMIM #158900) is one of the most prominent muscular dystrophies with an estimated prevalence of 1:8000 to 1:20000 [1,2]. FSHD is an autosomal dominant disorder that typically arise during the teenage years. Age-of-onset and severity vary, but all FSHD patients typically develop asymmetric muscle weakness in their facial, shoulder girdle and upper arm muscles, with progression to abdominal and lower limb muscles [3]. FSHD1 is caused by contraction of the D4Z4 repeat array to 1 – 10 repeat units (RUs) at the 4q35 locus, while non-FSHD individuals typically harbor more than 8 D4Z4 RUs [4]. The clinically indistinguishable FSHD2 type, carries normal D4Z4 RUs, but has the mutations in chromatin regulators such as SMCHD1, DNMT3B, and LRIF1 [5–7]. Whereas *DUX4* is normally transcriptionally silent in most somatic cells, disease manifestation in both types requires a disease-permissive 4qA haplotype containing a polyA signal (PAS) that stabilizes *DUX4* transcription [8–10].

Ectopically expressed *DUX4* can activate a cascade of genes unrelated to muscle functions, including early embryonic state-related genes [11], and cause cellular dysfunctions [12,13]. Therefore, suppressing *DUX4* transcription or translation is a potent therapeutic strategy to halt disease progression [14–17]. Currently, no cure or approved therapy for FSHD is available in the clinic [18,19].

While several reports have shown promising results in reducing *DUX4* activity through chemical approaches including RNA interference [20–24] or signal transduction inhibition [25], these strategies require lifelong continuous treatment because of chemical instability and degradation. Genome editing, however, may enable permanent transcriptional repression, but some reports have shown controversial results after removing the PAS on 4qA due to stochastic effects [26,27]. Targeting the *DUX4* coding sequence or promoter remains difficult because it resides in a repetitive D4Z4 unit. Moreover, genome editing strategies generally face technical and ethical issues because of irreversible off-target risks and the creation of unnatural mutations [28,29]. Beyond the distinct genotypes of FSHD, the D4Z4 array on the permissive allele is commonly epigenetically characterized by DNA hypomethylation in both FSHD1 and FSHD2, allowing abnormal chromatin relaxation and *DUX4* transcription in muscle cells [30]. Furthermore, large clinical cohort studies demonstrated that hypomethylation of the D4Z4 repeat unit on the 4qA haplotype is associated with disease severity and progression in FSHD [31,32]. Therefore, to prevent disease onset and progression, restoring the pathogenic D4Z4 DNA hypomethylation to a normal hypermethylated state that effectively blocks aberrant *DUX4* transcription represents a promising therapeutic strategy.

The CRISPR/Cas9 system, as a feasible platform for targeted genome editing, opened a new path for epigenome editing by creating a catalytically inactive or “dead” Cas9 (dCas9) with sustained sequence-specific targeting capacity [33–35]. Various tools for targeted epigenetic modulation based on dCas9 have been developed recently to regulate gene expression and thus represent potential therapeutic strategies for pathogenic conditions, including congenital diseases [36–38]. Indeed, it was reported that the Krüppel-associated box (KRAB) domain fused to dCas9 (dCas9-KRAB) and MeCP2 transcription repression domain (TRD) fused to dCas9 (dCas9-TRD) repressed *DUX4* expression in FSHD patient-derived myoblasts and an FSHD mouse model, respectively, indicating that targeting *DUX4* by dCas9-based epigenetic repression could be a promising strategy to cure FSHD [39,40]. As the KRAB domain, which mainly increases H3K9 tri-methylation via KAP1 recruitment, and the MeCP2 TRD, which primarily recruits histone deacetylase corepressor complexes, may only exert limited suppressive effects [41], stable and constitutive expression may be required for clinical application. However, a study showed that Cas9 protein induction triggered immune responses in a canine muscular dystrophy model [42], thus posing a potential barrier to constitutive expression of Cas9-associated components as a therapy.

Moreover, clinical trials testing AAV vectors—broadly considered a potent method for delivering transgenes to muscle tissue [43,44]—have led to patient deaths [45,46] and therefore have prompted questions about risks and benefits for FSHD patients with relatively mild and gradual disease progression in most cases. This problem could be avoided by developing an alternative method to sustain suppression of endogenous *DUX4* expression in patient cells by transient induction of dCas9 components with a suitable delivery method.

Among various modifications, DNA methylation is a more potent target to achieve sustained epigenetic editing effects when compared to histone modifications [41,47–49]. Moreover, it was reported that “hit-and-run”, combined induction of KRAB and DNA methyltransferases-associated domains, such as the DNMT3A catalytic domain, can induce lasting silencing effects at a targeted genomic site through the upregulation of local DNA methylation by transient transfection [50]. Moreover, the addition of enzymatically inactive DNMT3L can enhance DNA methyltransferase activity and “hit-and-run” effects [51–54]. Given that the chromatin state is tightly linked to DNA hypermethylation at the D4Z4 array in unaffected cells, in contrast to abnormal relaxation and DNA hypomethylation in FSHD cells, we considered targeting the D4Z4 array with a combination of these dCas9-mediated suppressive components to suppress endogenous *DUX4* expression. Here, using our validated patient-derived iPSC model with pathogenic levels of *DUX4* activity in the context of myogenic differentiation [55], we provide a proof of concept for the “hit-and-run” silencing strategy against endogenous *DUX4* gene expression in both FSHD1 and FSHD2, reversing disease-associated gene repression and pathway dysregulation, thereby highlighting potential clinical applications beyond distinct FSHD genetic backgrounds.

## MATERIAL AND METHODS

### Human iPSC clones and MyoD-induced myogenic differentiation

Human iPSC clones from a male FSHD1 patient, a female FSHD2 patient (for treatment with dCas9-effectors), a male FSHD2 patient (control)—specified as No Vector in the global methylation study—and one unaffected donor were originally described in a previous study [55]. Unaffected control S01 was generated with written consent from the volunteer and approval by the Kyoto University Graduate School and Faculty of Medicine Ethics Committee (approval nos. E1762, G567, and Rinsho71). iPSC clones with the doxycycline-inducible MyoD transgene (iPS^tet-MyoD^ clones) were established by transfecting them with plasmids, including *piggyBac* transposase (PBase) and PB-MyoD (with a puromycin resistance marker), and sub-cloning the clone with the highest differentiation efficiency. All human iPSC clones were cultured and maintained under feeder-free culture conditions as previously described [56]. Briefly, cells were passaged once a week and cultured on an iMatrix-511 (Nippi, Japan) pre-coated dish in StemFit AK02N medium (Ajinomoto). To select and maintain relevant transgenes, 0.5 μg/mL puromycin (Nacalai Tesque, Japan) or 100 μg/mL neomycin sulfate (Nacalai Tesque) was added. Muscle cell differentiation was performed as described previously [57] with some modifications. Briefly, at day 0, iPS^tet-MyoD^ cells were treated with Accutase (Nacalai Tesque) on day 0 to separate them into single cells and plated at a density of 2–2.5×10^5^ cells/well in 6-well dishes coated with Matrigel (Corning, NY) in StemFit supplemented with 10 μM Y-27632 (Nacalai Tesque). On day 1, the medium was replaced with primate ES cell medium (REPROCELL, Japan). On day 2, the culture medium was replaced with primate ES cell medium containing 1 μg/mL doxycycline (Dox; LKT Laboratories, MN). Induction was performed in 5% KSR/α-MEM containing Dox and 2-mercaptoethanol (2-ME) supplement without replating on day 3 for 3 days (FSHD1 clones) or 4 days (FSHD2 clones). After induction, cells were matured in 5% KSR/α-MEM containing 2-ME, 10 ng/mL human IGF-1 (Pepro Tech, UK), and 5μM SB431542 (Wako, Japan) until day 9 for FSHD1 clones and day 9 or 10 for FSHD2 clones.

### Vector construction and transfection

The dCas9-KRAB(BFP) vector (ID: VB190207-1015wke, VectorBuilder Japan) was originally designed and created using an online design tool (https://www.vectorbuilder.jp/design.html) with a *piggyBac* backbone and a dCas9-effector driven by a CAG promoter followed by an internal ribosome entry site (IRES) and blue fluorescent protein (TagBFP). The KRAB domain was sandwiched by EcoRI and AgeI restriction enzyme sites, which enabled the insertion of all other alternative effector domains including DNMT3A (D3A, amplified open reading frame, ORF, from Addgene #66819), DNMT3B (D3B, amplified ORF from Addgene #66820), DNMT3L (D3L, amplified ORF from Addgene #35523), and a negative control sequence. dCas9-D3A3L(BFP) vector was built by adding the D3L C-terminal domain ORF after the D3A domain on dCas9-D3A(BFP). Sequence ranges for each of the DNMT3A, DNMT3B, and DNMT3L domains were based on those in a previous report [58]. The dCas9-KRAB(Neo) vector was created by replacing the TagBFP of dCas9-KRAB(BFP) with a neomycin resistance gene, and dCas9-Neg(Neo) was built by replacing KRAB with a negative control sequence. The backbone of the single guide RNA (sgRNA) vectors was built by removing the hygromycin resistance gene from pPV-H1-ccdb-Ef1a-RiH_Ver1, provided by Dr. Akitsu Hotta (CiRA, Kyoto University, Japan), and sgRNA fragments were inserted as previously described [59]. Effector sequences and sgRNA target sequences are summarized in Supplementary Table S1 and S2, respectively. dCas9-effector and sgRNA vectors were transfected by a NEPA21 electroporator (Nepa Gene, Japan) into human iPS cells. A mixture of sgRNA vectors (2 µg total, comprising 500 ng of each of four sgRNA vectors), 12–13 µg of dCas9 vector, and 1 µg of PBase vector were co-transfected into 1 × 10^6^ cells in 100 µL of Opti-MEM (Thermo Fisher Scientific, Japan) per cuvette, as was described previously [48].

For the genuine hit-and-run system, an all-in-one vector CRISPRoff-v2.1(Addgene #167981) was used. Briefly, 4µL of Lipofectamine Stem Transfection Reagent (Invitrogen, cat. no. STEM00003) was mixed with 46 µL of Opti-MEM medium and then added to 50 µL of DNA solution containing 1 µg of total DNA (800 ng of the CRISPRoff vector and 200 ng of four sgRNA vectors). The resulting 100-µL transfection complex was incubated at room temperature for 10 min. iPSCs were dissociated using Accutase and resuspended in StemFit medium supplemented with 10 µM Y-27632. The cell concentration was adjusted to 5×10^5^ cells/mL. After incubation, the transfection complex was mixed with 1 mL of iPSC suspension, and the mixture was plated in a 12-well plate.

### Construction and delivery of CRISPRoff mRNA and sgRNA

CRISPRoff mRNA (ID:P230109-1018uaf, VectorBuilder Japan) used in this study was custom-synthesized via *in vitro* transcription. The template plasmid was derived from CRISPRoff-v2.1(Addgene #167981), where the BFP sequence was replaced by mCherry. The transcript contained a Cap1 structure and a poly(A) tail, and was purified by oligo (dT) affinity chromatography. N¹-methylpseudouridine (m¹Ψ) was incorporated to minimize immunogenicity and improve stability [60]. The sgRNA targeting the *DUX4* gene was also custom-synthesized by Custom Alt-R sgRNA service (ID:05840604Q, IDT Singapore). sgRNAs used in transient RNA delivery targeted the same sequences as those used in the plasmid-based transfection. To enhance sgRNA stability, 2′-O-methyl (2′-OMe) and phosphorothioate (PS) modifications were added to the three terminal nucleotides at both the 5′ and 3′ ends. Lipofectamine MessengerMAX Transfection Reagent (Invitrogen, cat. no. LMRNA003) was used to deliver CRISPRoff mRNA and sgRNA into iPSCs. First, 7.5 uL Lipofectamine MessengerMAX Transfection Reagent was mixed with 125 uL Opti-MEM and incubated for 10 min at room temperature. A 2.5 ug of RNA mixture (CRISPRoff mRNA 1.875 ug mixed with 0.625 ug of four sgRNAs) in 125 uL of Opti-MEM. The total CRISPRoff mRNA transfection complex (250 uL) was incubated at room temperature for 5 min. Finally, the transfection complex was added directly to iPSCs at 90% confluency, seeded in a 6-well plate one day before. For the experiment in iMuSCs, CRISPRoff mRNA and sgRNA amounts were reduced by half relative to those used in the iPSC model.

### Flow cytometry and cell sorting

Cells were dissociated with Accutase, washed once in StemFit medium containing 10 μM Y-27632, and resuspended in Hanks’ Balanced Salt Solution (HBSS) containing 10 μM Y-27632 and approximately 5% of the StemFit medium, then filtered and kept on ice. To monitor BFP (CRISPRoff) and RFP (sgRNA) fluorescence dynamics after transfection, fluorescence was measured using a BD LSRFortessa Cell Analyzer (BD Biosciences). To isolate fluorescent cells, we used a BD FACSAria II cell sorter (BD Biosciences). Cells transfected with dCas9-effector or CRISPRoff vectors with sgRNA vectors were sorted based on BFP and RFP fluorescence, while cells transfected with CRISPRoff mRNA were sorted based on mCherry fluorescence. Cells were then collected in culture medium with Y-27632 at 4℃ and replated for expansion. After expansion, cells were observed to capture phase-contrast and fluorescence images using a BZ9000 system (Keyence, Japan) at 200×magnification.

### RNA extraction and real-time reverse-transcription quantitative PCR (RT-qPCR)

Total RNA was extracted using a ReliaPrep RNA Miniprep System (Promega, WI) following the manufacturer’s instructions. cDNA was synthesized from the extracted RNA using a ReverTra Ace Master Mix with gDNA Remover (TOYOBO, Japan). Real-time RT-qPCR was performed using SYBR Green probe sets (Applied Biosystems, MA) and a Step One Plus thermal cycler (Applied Biosystems) or QuantStudio 3 (Applied Biosystems), and a standard curve was prepared for each target. RPLP0, which encodes a ribosomal protein, was used as the internal control in all assays. Primer sets used in this study are listed in Supplementary Table S3. Statistical analyses for all RT-qPCR were performed as described in the statistical analyses section.

### RNA-seq

Total RNA was obtained from sorted cells using the ReliaPrep RNA Cell Miniprep System (Promega). For Illumina sequencing libraries, 5 ng of total RNA was used with the SMART-Seq Total RNA-Seq Pico Input series (TAKARA, Japan) according to the manufacture’s instructions. The raw sequence data were converted to fastq files, and sequenced reads of R1 were trimmed to remove adaptors and low-quality bases using fastp (v0.24.0) with “-q 20 --length_required 100 --max_len1 100 --cut_front --cut_tail --cut_window_size 4 --cut_mean_quality 20” option. After filtering out reads that mapped to human ribosomal transcripts using Bowtie2 (v2.5.1), remaining R1 reads were mapped to the human reference genome GRCh38 using STAR(v2.7.11a) [61] with “--runModealignReads --winAnchorMultimapNmax1000 --outFilterMultimapNmax1000 --outMultimapperOrderRandom --outSAMmultNmax1 --alignSJoverhangMin8 --alignSJDBoverhangMin1 --outFilterMismatchNoverReadLmax0.04 --alignIntronMin 20 --alignIntronMa× 1000000 --alignMatesGapMa× 1000000” option. HTSeq(v2.0.5) [62] was used to calculate gene read counts, and the counted data were normalized by DESeq2(v1.42.0) [63]. Annotated gene lists representing *DUX4* target genes and myogenic differentiation-related genes were obtained from a previous study [64] and the Molecular Signatures Database (MSigDB; HALLMARK_MYOGENESIS, M5909) [65], respectively, and used for heatmap visualization. The corresponding gene lists are provided in Supplementary Table S4. To evaluate the effects of the CRISPRoff mRNA/sgRNAs system on the expression of *DUX4* target genes, Gene Set Enrichment Analysis (GSEA) was performed, and the expression patterns of leading-edge genes were visualized using heatmap. To assess global changes in gene expression, MA plots and GSEA were used to evaluate differentially expressed genes and enriched gene sets, respectively. For MA plot visualization, log2 fold changes were shrunk using the lfcShrink function to improve the stability of fold-change estimates, particularly for genes with low expression levels. In GSEA, the top 10 enriched gene sets ranked by normalized enrichment score (NES) and selected Hallmark pathways associated with FSHD pathogenesis and *DUX4*-related molecular processes were further examined. All RNA-seq data analyses and visualizations were performed using R.

### Targeted DNA methylation analysis using bisulfite sanger sequencing

Genomic DNA was extracted using a GenElute Mammalian Genomic DNA Miniprep Kit (Sigma-Aldrich, MO) according to the manufacturer’s protocol. A total of 500–1000ng of DNA was treated with bisulfite using an EpiTect DNA bisulfite kit (QIAGEN, Germany) according to the manufacturer’s guidelines. PCR and DNA methylation analysis of the *DUX4* 5′ promoter region for FSHD2 or 4qA region for FSHD1 was performed according to the protocol of the *DUX4* 5′ BSS assay or 4qA BSS assay, as described previously [66], with QUMA online software (http://quma.cdb.riken.jp/index_j.html) [67]. The analysis was performed with the following parameters: upper limit of unconverted CpHs = 5, lower limit of percent converted CpHs = 95.0, upper limit of alignment mismatches = 30, lower limit of percent identity = 90.0. The averaged DNA methylation percentages for each CpG site were summarized as heatmaps with GraphPad Prism 9.

### Global DNA methylation analysis with EPIC array data

The Infinium MethylationEPIC BeadChip Kit (Illumina) was used for methylation profiling of samples from the *piggyBac* system, while the Infinium MethylationEPIC v2.0 BeadChip Kit (Illumina) was employed for samples from the CRISPRoff mRNA/sgRNAs-based system (Fig.7A). All procedures were performed at the CiRA Foundation, Kyoto University, Japan, in accordance with the manufacturer’s protocol. First, a total of 500 ng genomic DNA was subjected to bisulfite conversion using the EZ DNA Methylation Kit (Zymo Research, CA, USA). Genome-wide DNA methylation levels were then assessed using either the Infinium MethylationEPIC BeadChip (Illumina) or the Infinium MethylationEPIC v2.0 BeadChip (Illumina), and arrays were scanned on an Illumina iScan System according to the manufacturer’s protocol. The extended EPIC RGset was generated from raw IDAT files.

Quality control (QC) was then performed, including assessment of control metrics (ewastools, v1.8.0), methylated and unmethylated signal intensities (minfi, v1.54.1), and sex prediction (ewastools, v1.8.0). Background correction using the noob method and functional normalization were subsequently performed with the minfi package. For CRISPRoff mRNA/sgRNAs datasets, the EPIC v2 annotation was updated to array = “IlluminaHumanMethylationEPICv2” and annotation = “20a1.hg38”, whereas *piggyBac* system datasets were processed using the original EPIC v1 annotation (array = “IlluminaHumanMethylationEPIC” and annotation = “ilm10b5.hg38”). Given that EPIC v2 contains 5,225 probes with two or more replicate measurements, replicate probes were collapsed based on detection P values, with the replicate showing the lowest detection P value retained as the representative probe to maximize compatibility with EPIC v1 [68]. The 721,347 probes shared between EPIC v1 and EPIC v2 were retained for downstream analysis of all samples. Differentially methylated probes (DMPs) were identified using the limma package, with a β-value difference > 0.20 and an adjusted P-value < 0.01 as the significance thresholds. Genes harboring more than three DMPs were selected for subsequent analysis and were listed in Supplementary Table S5. To integrate EPIC datasets and RNA-seq data, the EnhancedVolcano package (Blighe, Rana, and Lewis, 2018) was used to generate volcano plots, with selected genes highlighted. Gene labels were displayed only for highlighted genes meeting the significance criteria, defined as an absolute log2fold change (|log2FC|) ≥ 1 and a P-value ≤ 1×10^-5^. Principal component analysis (PCA) was performed in R using DNA methylation β-values from the common probes.

### In silico off-target prediction

The 23-bp target sequence of each sgRNA (20-bp + 3-bp PAM) was submitted to the CRISPRdirect online tool [69] to identify potential off-target sites matching the 20-mer, 12-mer, and 8-mer sequences upstream of the PAM (NGG), together with their genomic coordinates. Since EPIC v1 array annotation is based on hg19 in the CRISPRdirect online tool [69], the off-target prediction sites of tKD vs. tNN were obtained using the hg19 assembly. In contrast, off-target sites for the LP^3^ vs. NT comparison were identified using the hg38 assembly to match the EPIC v2 array annotation. Candidate off-target sites matching either the 8-mer or 12-mer sequence upstream of the PAM were retained for subsequent analysis. The Illumina HumanMethylationEPIC anno.ilm10b5.hg38 annotation package (EPIC v1), an extended version of hg19 for more compatible analysis with EPIC v2 array, and the Illumina HumanMethylationEPIC v2 anno.20a1.hg38 annotation package (EPIC v2) were loaded in R. Probes retained after quality filtering were converted to genomic ranges with gene annotations. Candidate methylation probes located within 500 bp of each predicted off-target site were identified using the ‘findOverlaps()’ function and filtered based on a maximum distance of 500 bp. This in silico workflow was confirmed using a newly designed sgRNA target against the MyoG promoter as a virtual example, as it successfully identified the MyoG gene and other potential off-target genes (Supplementary Fig. S7E). The off-target sites of the four sgRNAs targeting *DUX4*-fl were identified using this in silico workflow, and the gene annotation list for each sgRNA is shown in Supplementary Table.S6. The code for the prediction process is available in GitHub (https://github.com/mits2112/offtarget_predict_EPICarray).

### Differentiation of iPSCs into iMuSCs and myotubes

A stepwise differentiation protocol developed in our laboratory was used [70]. iPSCs were seeded at a density of 2×10^3^ to 2×10^4^ cells per well in Matrigel-coated 24-well plates (BD Biosciences) and cultured in StemFit (AK02N, Ajinomoto), supplemented with 10 µM ROCK inhibitor Y-27632 (Sigma), for 3 days. First, dermomyotome-like cells were induced over 14 days using differentiation medium consisting of CDMi supplemented with CHIR99021 (CHIR; Axon MedChem, Tocris) and SB431542 (SB; Sigma). On day 14, cells were passaged and cultured in CDMi supplemented with Y-27632 (10 µM) for 3 days. Myogenic differentiation was then initiated on day 17 by switching to serum-free culture medium (SF-O3; Sanko Junyaku) supplemented with 0.2% bovine serum albumin (BSA), 0.1 mM 2-mercaptoethanol (2-ME), 10 ng/mL recombinant human IGF-1 (PeproTech), 10 ng/mL recombinant human bFGF (Oriental Yeast Co., Ltd.; NIB 47079000), and 10 ng/mL recombinant human HGF (PeproTech). From day 38 onward, cells were cultured in muscle maturation medium consisting of DMEM (Invitrogen, 11960069) supplemented with 0.5% penicillin–streptomycin (Nacalai, 26253-84), 2 mM L-glutamine (Nacalai, 16948-04), 0.1 mM 2-ME, 2% horse serum (HS; Sigma), 5 µM SB431542, and 10 ng/mL IGF-1 (PeproTech) until week 12. To isolate skeletal muscle stem cells, FACS sorting was conducted based on surface markers CD82+CD57-[71] or the more distinct iMuSC marker CDH13[72]. Sorted iMuSCs were seeded at 3×10³ to 5×10³ cells per well in Matrigel-coated 96-well plates. iMuSCs were cultured in StemFit until confluence after approximately 1 week, followed by myogenic differentiation medium consisting of DMEM supplemented with 1% N2 (Thermo Fisher Scientific).

### Statistical Analyses

For RT-qPCR data from dCas9-effector vector experiments, statistical analysis was performed using one-way ANOVA, followed by Tukey’s multiple comparisons test in comparisons among three or more groups. Unpaired Student’s t-test was used for the comparison between two independent groups. For RT-qPCR analysis of *SLC34A2* and *CCNA1*, two-way ANOVA followed by Tukey’s multiple comparisons test was used. All statistical tests were two-sided. All RT-qPCR data were processed using GraphPad Prism 9 for visualization and statistical analysis and were represented as mean±SEM. *P < 0.05, **P < 0.01, ***P < 0.001, ****P < 0.0001, and ns indicates not significant. Fisher’s exact test was used to compare the proportions of hypermethylated and hypomethylated probes between tKD vs. tNN groups and LP3 vs. NT groups using R. Further details are provided in the figure legends.

### Ethical Approval

This study was approved by the Ethics Committee of the Graduate School of Medicine, Kyoto University, and the Kyoto University Hospital (Approval #R0091 and #G259) and was conducted according to the guidelines of the Declaration of Helsinki. To protect confidentiality, all patient information has been kept anonymous, with written informed consent was obtained from the study participants.

## RESULTS

### Development and validation of dCas9-effector activity

First, we constructed *piggyBac*-based vectors expressing dCas9-mediated repressive effectors and sgRNAs targeting the D4Z4 array, using *piggyBac* transposase (PBase) enabling switching between transient and constitutive expression of these components (Supplementary Fig. S1A). Effectors included KRAB, catalytic domains of DNMT3A (D3A) and DNMT3B (D3B), a pseudo-catalytic domain of DNMT3L (D3L), an enhanced D3A–D3L fusion effector (D3A3L), and a negative control sequence (Neg). Four sgRNAs targeting the promoter region of the D4Z4 array were selected from validated sgRNAs reported by Himeda et al. with dCas9-KRAB [39]. Each dCas9-effector vector was electroporated together with four sgRNA vectors into FSHD2 iPS^tet-MyoD^ cells (FSHD2 iPSCs), which had previously been established as an FSHD patient-derived iPSC model exhibiting robust expression of *DUX4* and its downstream targets following myogenic differentiation [55] (Fig. 1A). One week after electroporation, BFP- and RFP-double-positive cells were sorted and cultured for an additional week to allow expansion, after which they were differentiated into myocytes (Supplementary Fig. S1B). *DUX4* and its downstream targets (*ZSCAN4*, *MBD3L2*, and *TRIM43*) were significantly suppressed by dCas9-KRAB, consistent with a previous report (Fig. 1B, C). dCas9-D3A, dCas9-D3L, and dCas9-D3A3L, but not dCas9-D3B, were able to suppress *DUX4* expression. However, suppression was lower than that achieved by dCas9-KRAB. Myogenic differentiation was comparable across all conditions (Fig. 1D). DNA methylation analysis conducted by bisulfite sequencing (BSS) revealed widespread increases in DNA methylation across the D4Z4 repeat unit after treatment with dCas9-KRAB, dCas9-D3A, and dCas9-D3AD3L, whereas dCas9-D3L produced only partial methylation (Fig. 1E, Supplementary Fig. S1C). These data demonstrated the enzymatic activity of the dCas9-effectors and confirmed that our FSHD-iPSC model is suitable for evaluating their *DUX4*-suppressive effects.

**Figure 1:**
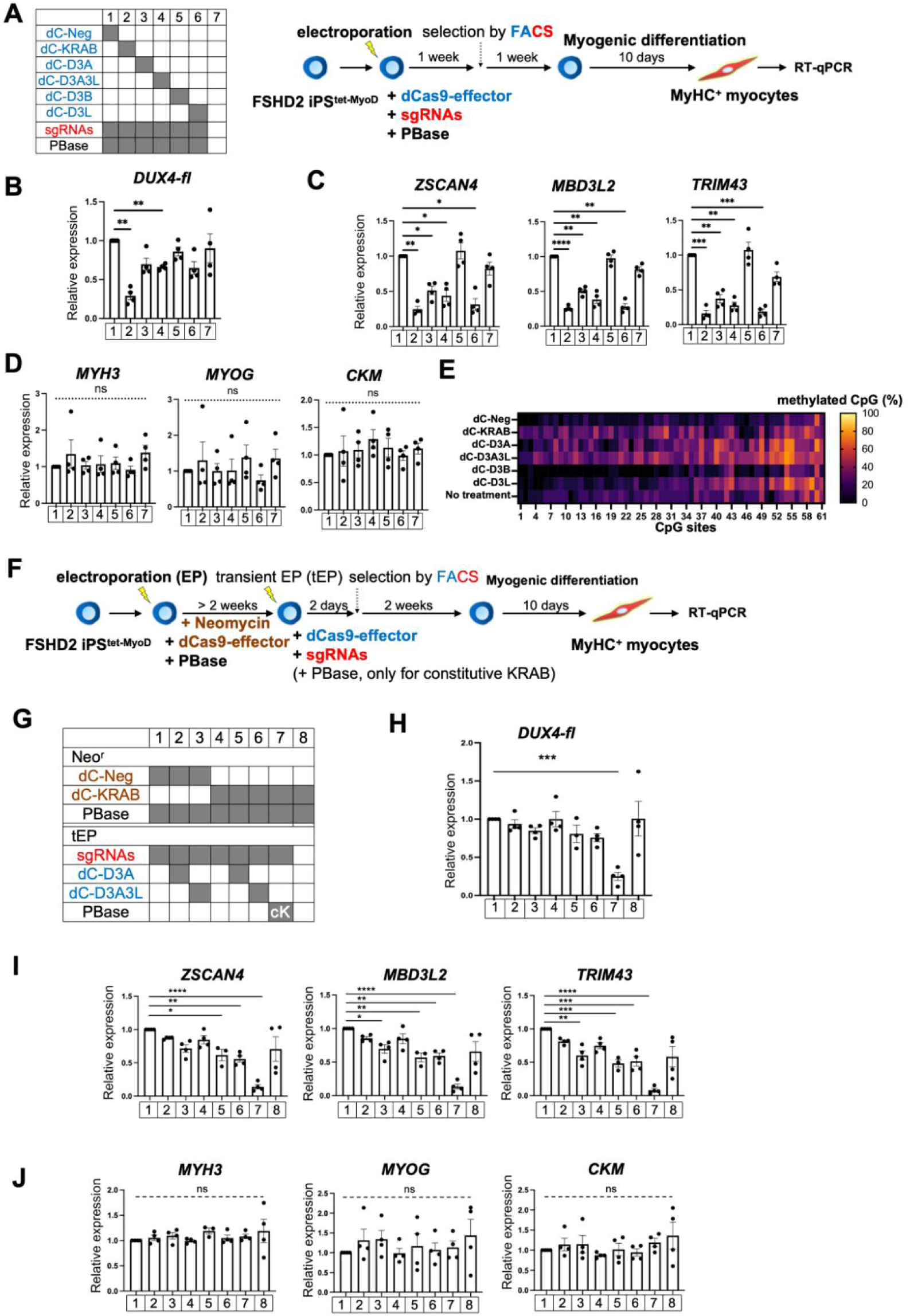
Establishment and validation of dCas9-effectors for the evaluation of *DUX4* suppression, revealing that a combination of dCas9-KRAB and dCas9-D3A/D3A3L suppressed *DUX4* with transient transfection. See also Supplementary Fig. S1. A) Scheme showing the transfection, selection, and differentiation of iPSC clones to myocytes. After FSHD2 iPSC^tet-MyoD^ clones were transfected by electroporation, cells with constitutive BFP and RFP dual signals were sorted and differentiated for analysis. B-C) RT-qPCR analysis for B) *DUX4-fl*, C) *DUX4* downstream targets, and D) a myogenic differentiation marker in cells on day 10 of differentiation. Relative expression was normalized to dC-Neg in each experimental set (n = 4). E) DNA methylation analysis of the D4Z4 regions using bisulfite sequencing. Note that all the sgRNA targets are located inside the D4Z4 unit but outside of the analyzed region. F) Scheme showing the transfection, selection, and differentiation of iPSC clones to myocytes. FSHD2 iPSC^tet-MyoD^ clones were co-transfected by electroporation with the dCas9-effector(Neo) and PBase together for constitutive expression, then grown for > 2 passages to remove PBase. Cells were co-transfected by electroporation with dCas9-effector(BFP) and sgRNAs(RFP) together for transient expression. After two days, cells with BFP and RFP dual signals were sorted, expanded for two weeks, and differentiated for analysis. Note that the dCas9-effector is targeted only when sgRNAs are induced in cells, except under condition 7, where PBase was added again in the second electroporation to express sgRNAs constitutively. G) Experimental design for comparison. While dCas9-KRAB was expressed continuously, transient expression of sgRNAs enabled the transient targeting of dCas9-effectors at target site since PBase was excluded for 1–6. In addition, dCas9-KRAB(Neo) transfected with sgRNAs and PBase (7. cK) was also prepared as a positive control. H-J) RT-qPCR analysis for H) *DUX4-fl*, I) *DUX4* downstream targets, and J) a myogenic differentiation marker in cells on day 10 of differentiation. Relative expression was normalized to condition 1 in each experimental set (n = 4). All RT-qPCR data are represented as mean ± SEM. *P < 0.05, **P < 0.01, ***P < 0.001, ****P < 0.0001, and ns indicates not significant as determined using one-way ANOVA followed by the Tukey’s multiple comparisons test.

### Synergistic effect of dCas9-KRAB and dCas9-D3A/D3A3L

Next, prompted by the observed partial *DUX4* suppression and D4Z4 methylation, and by previous reports that transient delivery of a single effector has limited suppression effect [50], we evaluated whether a combination of dCas9-KRAB and dCas9-D3A/D3A3L by transient delivery could achieve a hit-and-run effect. Since dCas9 constructs are too large to be co-delivered into individual cells after a single transfection, the system was modified with the dCas9-KRAB construct carrying a neomycin resistance gene [dCas9-KRAB(Neo)], instead of BFP (Supplementary Fig. S1D). This meant that selected cells expressed both dCas9-KRAB(Neo) and dCas9-D3A/D3A3L(BFP), following selection by neomycin resistance and FACS based on BFP fluorescence. First, FSHD2 iPSCs were co-transfected with dCas9-KRAB(Neo) or dCas9-Neg(Neo) and PBase to achieve stable genomic integration and constitutive expression, followed by maintenance under neomycin selection (Fig. 1F). Then, Cells were transiently electroporated with (i) dCas9-D3A plus four sgRNAs, (ii) dCas9-D3A3L plus four sgRNAs, or (iii) four sgRNAs alone. In addition, constitutive induction of KRAB and four sgRNAs (cK) was prepared as a positive control of *DUX4* suppression (Fig. 1G). Two days after transfecting dCas9-D3A/D3A3L(BFP) and sgRNAs, cells were sorted for dual RFP^+^BFP^+^ positive cells, and myogenic differentiation was conducted following two weeks of cell expansion (Fig. 1F, Supplementary Fig. S1E). Gene expression analysis showed that transient activation of dCas9-KRAB alone failed to suppress *DUX4.* Whereas transient activation of dCas9-KRAB with either dCas9-D3A or dCas9-D3A3L achieved modest suppression of *DUX4* and its target genes, the effect did not reach the level observed with cK (Fig. 1H, I). Myogenic differentiation was not significantly affected under any condition, indicating that the modest suppression of *DUX4* observed in the combination group can be attributable to activities held on dCas9-effectors rather than impaired muscle differentiation (Fig. 1J). These data indicated that the combined activity of dCas9-KRAB and dCas9-D3A/-D3A3L, but not dCas9-KRAB alone, resulted in modest suppression of *DUX4* even after two weeks of expansion and 10 days of myogenic differentiation.

### Repeated treatment enhanced the suppression effect in the FSHD2 model

Since a slight but significant effect on *DUX4* targets was observed following transient activity of the combination (Fig. 1I), we hypothesized that repeated transfection would enhance the effect. Accordingly, dCas9-KRAB(Neo) cells underwent one, two, and three rounds of transfection with dCas9-D3A3L and sgRNAs at 7-day intervals (Fig. 2A, B). Two days after each transfection, RFP- and BFP-double-positive cells were sorted by FACS (Supplementary Fig. S2A–C). One week after electroporation, most cells were negative for both BFP and RFP (Supplementary Fig. S2D). Cells were then expanded for an additional week before differentiation into myocytes. The expression of *DUX4* and its downstream targets was analyzed in differentiated myocytes. While the transient activity of dCas9-KRAB(Neo) alone failed to suppress *DUX4*, the combination of dCas9-KRAB (Neo) and dCas9-D3A3L resulted in progressively stronger *DUX4* suppression after one (tEP^1^), two (tEP^2^), and three (tEP^3^) rounds of transfection, without affecting myogenic differentiation (Fig.2C–E). Notably, suppression of *DUX4* and its target genes after three rounds of transfection was comparable to that observed in cK cells, which constitutively express dCas9-KRAB and sgRNAs (Fig. 2C, D). Moreover, RNA-seq analysis of tEP^3^ cells confirmed widespread repression of *DUX4* target genes without compromising myogenic differentiation (Fig. 2F-H, Supplementary Table S4). Consistent with previous clinical genetic studies [66], our FSHD2 iPSC model exhibited D4Z4 hypomethylation compared with unaffected control cells. Following tEP^3^ treatment, DNA methylation was dramatically increased over the entire analyzed region of the D4Z4 array (Fig. 2I, Supplementary Fig. S2E). These data suggest that repeated induction of combined activity of dCas9-KRAB and dCas9-D3A3L establishes sustained *DUX4* suppression in FSHD2 cells through robust D4Z4 DNA hypermethylation.

**Figure 2.**
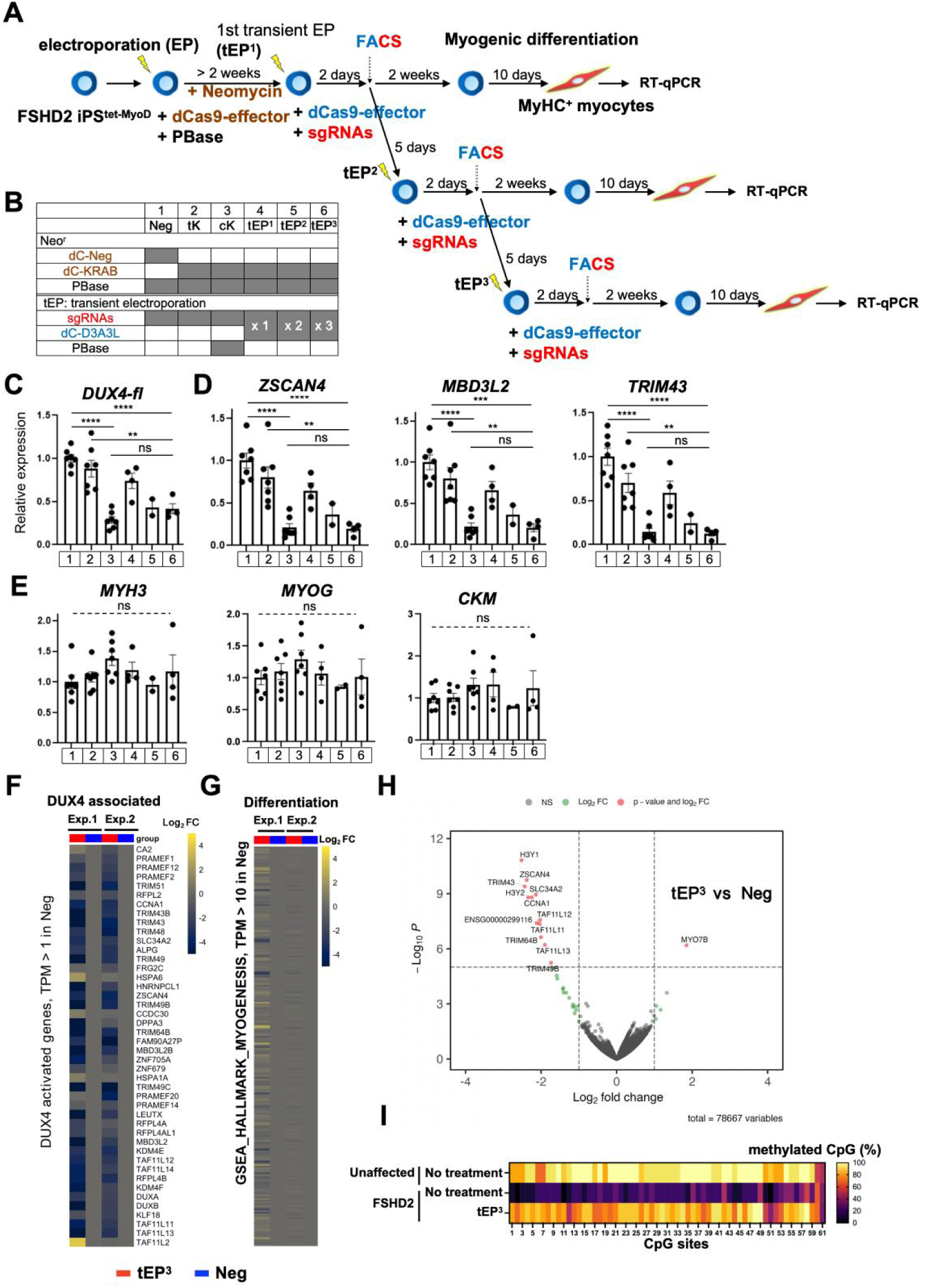
Repeated transfection boosted the silencing effect of transient combinations. See also Supplementary Fig. S2. A) Scheme showing the transfection, selection, and differentiation of iPSC clones to myocytes. FSHD2 iPSC^tet-MyoD^ clones with constitutive dCas9-effector(Neo) were transfected with dCas9-effector (BFP) and sgRNAs by electroporation for transient expression (the first transient electroporation, tEP^1^). On day 2 after transfection, cells were sorted with BFP and RFP dual signals. After 2 weeks of cell expansion, some of sorted cells were subjected to MyoD-induced myogenic differentiation for *DUX4*-associated gene expression analysis by tEP^1^. In addition, some sorted tEP^1^ cells were subjected to a second round of transfection (tEP^2^). The process was similarly applied to cells sorted after tEP^2^ for BFP and RFP dual signals, with tEP^2^ cells differentiated for analysis after two weeks of expansion. Furthermore, some of tEP^2^ cells were subjected to a third round of transfection (tEP^3^), and analyzed after the same sorting, expansion, and differentiation process. Note that the dCas9-effector is targeted only when sgRNAs are induced in cells, except under condition 3, where PBase was added again for constitutive sgRNA expression. B) Experimental design for comparison. dCas9-KRAB (Neo) integrated cells are transiently transfected with sgRNAs and dCas9-D3A3L (4-6. tEP^1^, tEP^2^ and tEP^3^).

In addition, dCas9-Neg(Neo) transfected with sgRNAs as a negative control (1. Neg), and dCas9-KRAB(Neo) transfected with sgRNAs without PBase (2. transient KRAB induction, tK), and with PBase as a positive control (3. constitutive KRAB induction cK) were also prepared. C-E) RT-qPCR analysis for C) *DUX4*-fl, D) *DUX4* downstream targets, and E) a myogenic differentiation marker in cells at day 10 of differentiation. Relative expression was normalized to the average for condition 1 in all samples. (n = 2–7). All data are represented as mean ± SEM. *P < 0.05, **P < 0.01, ***P < 0.001, ****P < 0.0001, and ns indicates not significant, as determined using one-way ANOVA followed by the Tukey’s multiple comparisons test. F-G) Heatmaps showing the gene expression determined from RNA-seq analysis of the differentiated Neg and tEP^3^ cells from two independent experimental sets F) for the *DUX4-*associated gene set and G) the myogenic differentiation-associated gene set. Exp.1 and Exp.2 indicate biologically independent experimental sets. H) Volcano plot generated using the gene expression data from the RNA-seq analysis for all genes. Note that *DUX4* target genes are significantly downregulated while most of genes are not significantly changed (-log_10_P≤ 5). I) DNA methylation analysis for the D4Z4 regions determined by bisulfite sequencing. For the non-treated FSHD2 clone, the same data as in Fig. 1E were used.

### Hit-and-run silencing could also suppress *DUX4*in the FSHD1 model

Encouraged by the successful suppression of *DUX4* by restoring of DNA methylation at the D4Z4 array in FSHD2 cells, we next examined whether the same hit-and-run strategy could also be applied to FSHD1 cells—characterized by D4Z4 repeat contraction-associated hypomethylation on one 4qA allele harboring fewer D4Z4 repeat units than in FSHD2—to investigate its broader applicability across FSHD subtypes. Because the electroporation efficiency of our FSHD1 iPS^tet-MyoD^ clone was low, we first established FSHD1 iPS^tet-MyoD^ cells (FSHD1 iPSCs) stably expressing dCas9-KRAB (Neo) and dCas9-DNMT3A3L (BFP) using the *piggyBac* (PBase) transposon system, followed by drug selection and FACS to isolate double-positive cells (Fig. 3A, B). These cells were subsequently subjected to three rounds of transient transfection with four sgRNAs (RFP), without PBase, to direct the dCas9-effector to its intended target. To isolate RFP-positive cells, FACS was performed using the RFP⁺ /BFP⁺ gate two days after each round of transfection. After the third sort, the resulting cells were expanded for two weeks followed by myogenic differentiation. To first validate this approach, FSHD2 cells subjected to this protocol showed *DUX4* suppression and increased DNA methylation at the D4Z4 region, consistent with our previous results (Supplementary Fig. S3). RFP fluorescence was no longer detectable one week after the third round of transfection, indicating loss of sgRNA vectors and the transient nature of dCas9-effector targeting activity (Fig. 3C, Supplementary Fig. S3C). We applied this approach to investigate the hit-and-run effect in FSHD1. Notably, hit-and-run treatment significantly suppressed *DUX4* and its downstream targets (Fig. 3D–F). While transient dCas9-KRAB activity (tKN) failed to repress *DUX4*, the transient combined activity of dCas9-KRAB and dCas9-D3A3L (tKD) achieved robust *DUX4* silencing, comparable to that achieved by constitutive dCas9-KRAB expression (cK) (Fig. 3B,D,E). Collectively, these results suggest that transient combined activity of dCas9-KRAB and dCas9-D3A3L is sufficient to achieve robust and sustained suppression of *DUX4* and its downstream targets in both FSHD1 and FSHD2 cells, providing a rationale to overcome the dCas9-effector packaging issue and to test whether a genuine hit-and-run silencing via transient delivery of all components is sufficient to achieve sustainable *DUX4* silencing.

**Figure 3.**
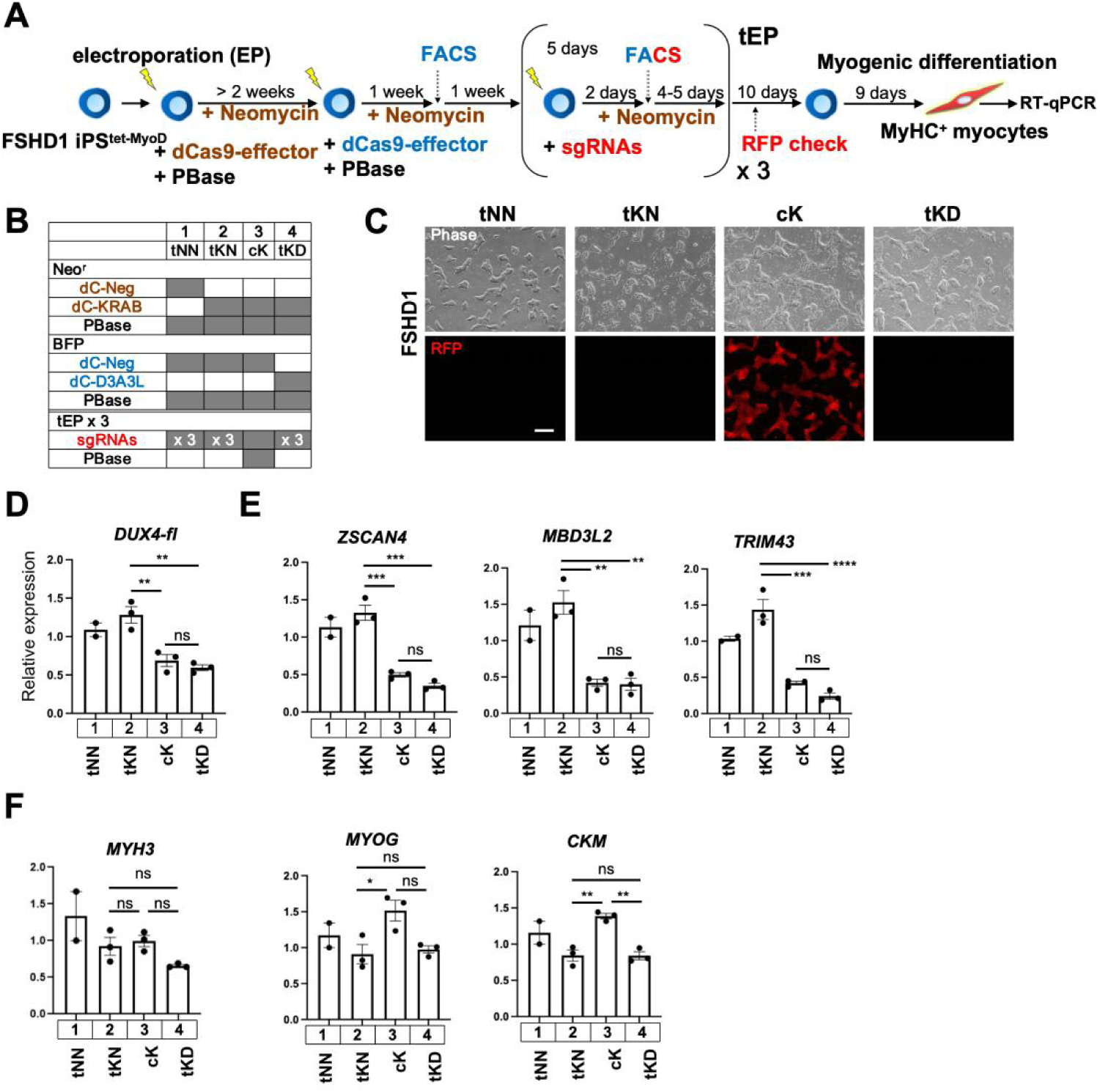
*DUX4* was suppressed by the transient activity of the dCas9-KRAB and dCas9-D3A3L combination in the FSHD1-iPSC model. See also Supplementary Fig. S3. A) Scheme showing the transfection, selection, and differentiation of FSHD1 iPSC clones to myocytes. FSHD1 iPSC^tet-MyoD^ clones were electroporated with a combination of dCas9-effector(Neo) and PBase for constitutive expression, then co-transfected with dCas9-effector (BFP) and PBase for constitutive expression and sorted based on BFP signal. After > 2 passages to remove the PBase after electroporation, sgRNAs were induced transiently and after two days sorted for BFP and RFP dual signals and expanded for 4 or 5 days. The transfection-sorting round was repeated three times, followed by a two-week expansion period and muscle differentiation. Please note that the dCas9-effector is targeted only when sgRNAs are induced in cells, except under condition 3, where PBase was added again in the second electroporation for constitutive sgRNA expression. B) Experimental design for comparison analysis. Sample conditions were named as tNN (transient sgRNAs with dCas9-Neg(Neo) and dCas9-Neg(BFP)), tKN (transient sgRNAs with dCas9-KRAB(Neo) and dCas9-Neg(BFP)), cK (constitutive sgRNAs with dCas9-KRAB(Neo) and dCas9-Neg(BFP)), and tKD (transient sgRNAs with dCas9-KRAB(Neo) and dCas9-D3A3L(BFP)). C) Phase-contrast and fluorescence images of RFP one week after the third round of transfection. Scale bar: 500 µm. D – F) RT-qPCR analysis for D) *DUX4-fl*, E) *DUX4* downstream targets, and F) a myogenic differentiation marker in cells on day 9 of differentiation. Relative expression was normalized to one sample from condition 1 [n = 3, except for condition tNN (n = 2)]. All data are represented as mean ± SEM. *P < 0.05, **P < 0.01, ***P < 0.001, ****P < 0.0001, and ns indicates not significant as determined using one-way ANOVA followed by the Tukey’s multiple comparisons test.

### Genuine hit-and-run system using all-in-one CRISPRoff vector

Efficient co-delivery of multiple dCas9-effectors remains a key obstacle to the clinical translation of epigenome editing. The all-in-one CRISPRoff vector developed by Weissman JS et al. [53], which encodes both dCas9-KRAB and DNMT3A/3L, enabled us to test a genuinely transient transfection. FSHD2 iPSCs underwent lipofection with the CRISPRoff (BFP) vector and four sgRNAs (RFP) vectors. RFP- and BFP-double-positive cells were sorted on day 2 after the lipofection (Fig. 4A, Supplementary Fig. S4A) followed by a second and third round of lipofection, and sorting for BFP^+^/RFP^+^ cells. Analysis of BFP and RFP signals on day 7 after each transfection round showed that the proportion of BFP^+^/RFP^+^ cells decreased to nearly zero. After an additional week of expansion, cells became completely negative for both BFP and RFP signals by day 13, immediately before differentiation (Fig. 4C). To further validate the genuine hit-and-run transfection, sorted BFP+/RFP+ cells were monitored daily for two weeks after the first transfection. The CRISPRoff-BFP signal became undetectable by day 5 post-transfection, and the sgRNA-RFP signal disappeared completely by day 10, indicating that both CRISPRoff vector and sgRNA vectors were transiently transfected without genomic integration (Fig. 4C,D). Whereas cells treated with CRISPRoff alone (Neg^1^, Neg^2^, and Neg^3^) showed no *DUX4* suppression, cells subjected to one, two, and three rounds of CRISPRoff and sgRNA transfection (HAR^1^, HAR^2^, and HAR^3^) exhibited progressive *DUX4* suppression, indicating that *DUX4* suppression was mediated by CRISPRoff-induced epigenetic editing rather than by the CRISPRoff protein itself (Fig. 4E – G). Moreover, DNA methylation on the D4Z4 array was broadly increased in HAR^3^ cells when compared to the original FSHD2 iPSCs (Fig. 4H, Supplementary Fig. S4B). These data demonstrated that the genuine hit-and-run delivery of CRISPRoff and sgRNAs achieved sustained *DUX4* suppression in FSHD2 cells through DNA hypermethylation, thereby addressing the challenge of packaging and transiently delivering multiple dCas9-effectors into cells.

**Figure 4.**
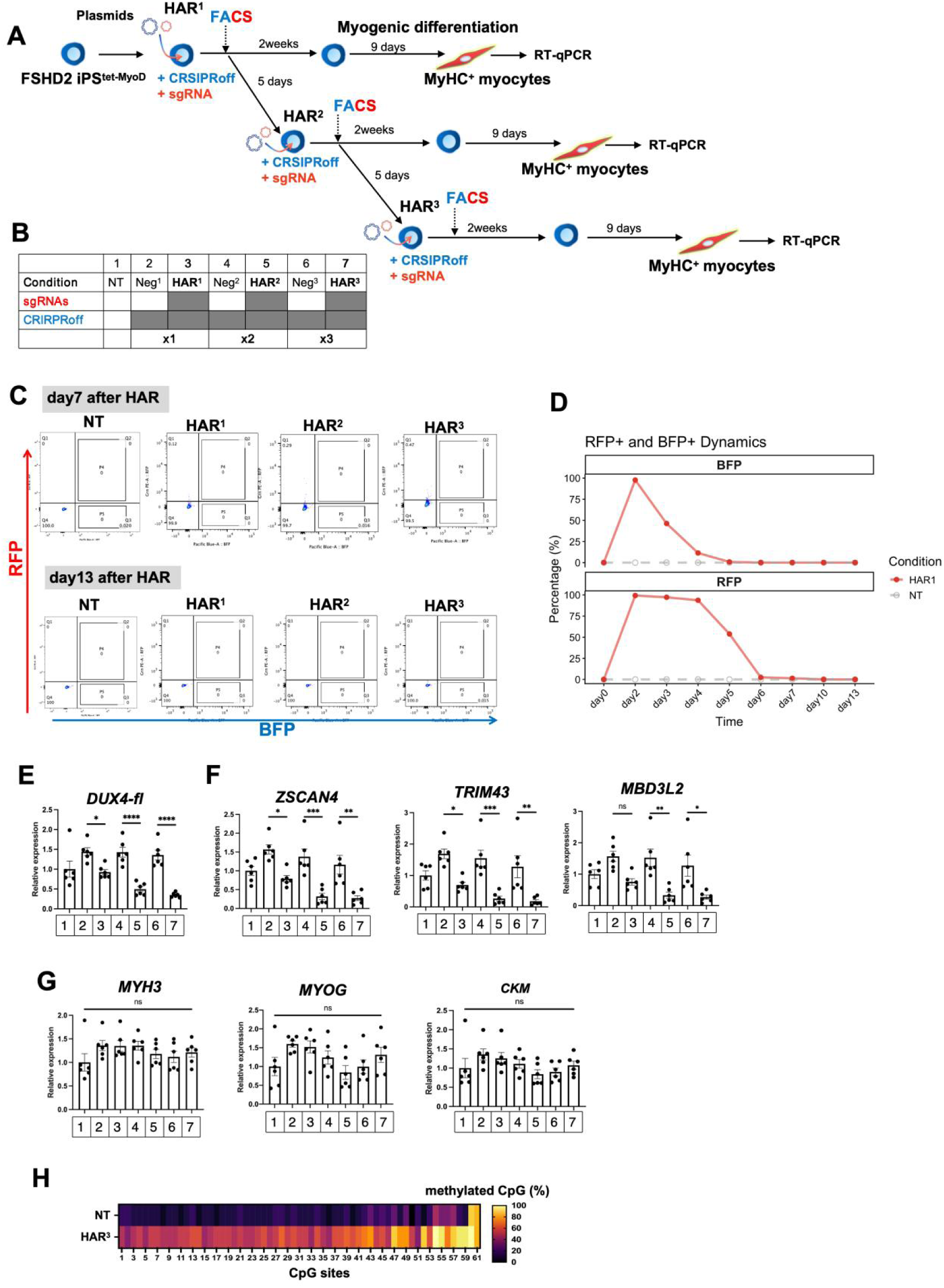
Using the all-in-one CRISPRoff vector, a genuine transient transfection system was successfully established for frequency-dependent *DUX4* suppression. See also Supplementary Fig. S4. A) Scheme showing the transfection, selection, and differentiation of FSHD2 iPSC clones to myocytes. FSHD2 iPSCs^tet-MyoD^ clones were transfected by lipofection (HAR^1^) with CRISPRoff (BFP) and sgRNAs (RFP) without PBase for transient expression. Two days later, cells with BFP and RFP dual signal were sorted. After a two-week expansion period, cells were differentiated into myocytes for analysis. In addtion, some HAR^1^ cells were subjected to a second round of transfection (HAR^2^). After the second transfection, cells were sorted for BFP and RFP dual signal. Analysis was performed two weeks after myocyte differentiation. A third round of transfection, sorting, cell expansion and differentiation was also conducted (HAR^3^) following the same process as the first and second rounds. B) Experimental design for comparison. FSHD2 iPSCs with one, two, and three rounds of CRISPRoff and sgRNA together(conditions 3, 5, 7: HAR^1^, HAR^2^, and HAR^3^). In addition, FSHD2 iPSCs were transfected with one, two, and three rounds of CRISPRoff only as negative controls (conditions 2, 4, 6: Neg^1^, Neg^2^, Neg^3^). C) BFP and RFP fluorescence on days 7 and 13 post-transfection were measured by flow cytometry. On day 7, almost all cells were negative. By day 13, before differentiation, plasmid-derived fluorescence signals became undetectable. D) Dynamics of RFP (sgRNA) and BFP (CRISPRoff) signals following HAR^1^. The BFP signal decreased to near background levels by approximately day 5 post-transfection, while the RFP signal remained detectable until around day 7. E) *DUX4*-fl, F) *DUX4* downstream targets, and G) myogenic differentiation markers in cells on day 9 of differentiation. Relative expression was normalized to the average for condition 1 in all samples (n=6). All data are represented as mean ± SEM. *P < 0.05, **P < 0.01, ***P < 0.001, ****P < 0.0001, and ns indicates not significant as determined using one-way ANOVA followed by the Tukey’s multiple comparisons test. H) DNA methylation analysis for the D4Z4 regions determined by bisulfite sequencing.

### The induction of CRISPRoff mRNA/sgRNAs achieved *DUX4*silencing in both FSHD2 and FSHD1

As the hit-and-run system leverages transient gene-delivery modalities, several studies have reported the successful use of RNA-based epigenetic editors via non-viral delivery at a single genomic locus [73–75]. Here, we evaluated whether RNA can deliver an epigenetic editor into FSHD cells to target the D4Z4 repeat array. CRISPRoff mRNA was synthesized in which BFP was replaced with mCherry to detect transfected cells, while the sgRNAs carried no fluorescent label (Supplementary Fig. S5A). CRISPRoff mRNA and sgRNAs were transfected into FSHD2 iPSCs with three transfections were performed at 7-day intervals. mCherry-positive cells were isolated by FACS 24h after each transfection. After isolating mCherry-positive cells following one, two, or three transfections, cells were expanded for two weeks and subsequently subjected to myogenic differentiation (Fig. 5A, B, Supplementary Fig. S5B). *DUX4* suppression was observed and enhanced by repeated transfection of CRISPRoff mRNA/sgRNAs (LP^1^, LP^2^, LP^3^) without impairing myogenic differentiation (Fig. 5C – E). Notably, the DNA methylation at the D4Z4 promoter region was robustly increased after the third transfection, comparable to that of the unaffected control (Fig. 5F, Supplementary Fig. S5C). These results revealed that transient delivery of CRISPRoff mRNA and sgRNAs can achieve stably suppression of *DUX4* and downstream targets, indicating that a hit-and-run CRISPRoff mRNA/sgRNAs approach efficiently induces epigenetic silencing at the D4Z4 repeat array.

**Figure 5.**
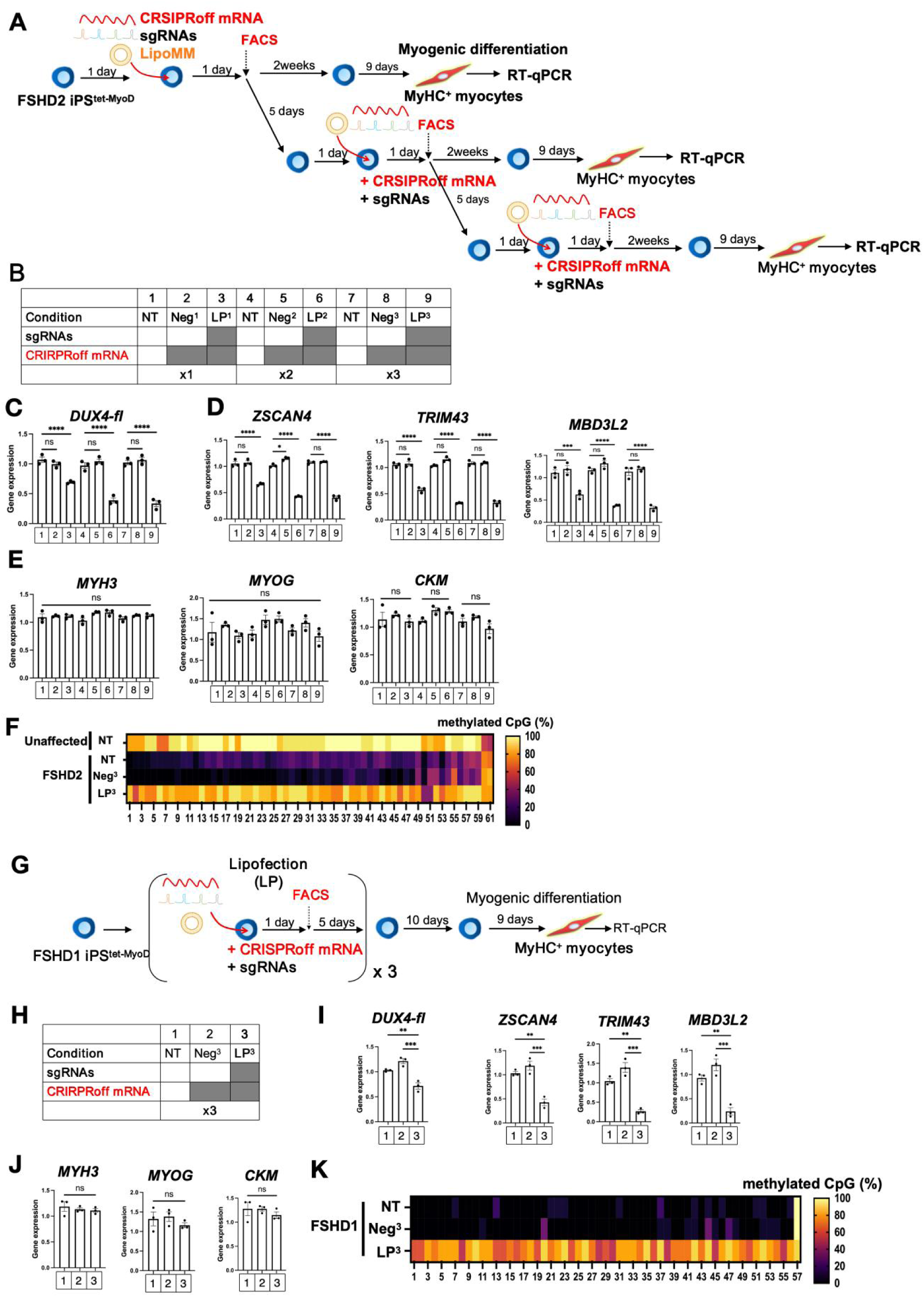
Transient transfection of CRISPRoff mRNA and sgRNA enables *DUX4* silencing in both FSHD2 and FSHD1 cells. See also Supplementary Fig. S5. A) Schematic of the transfection, selection, and differentiation workflow for FSHD2 iPSC clones into myocytes. FSHD2 iPSC^tet-MyoD^ clones were transfected by lipofection (LP^1^) with CRISPRoff mRNA (mCherry) and sgRNAs for transient expression. One day after seeding, mCherry-positive cells were sorted. After two weeks of expansion, myogenic differentiation was initiated for analysis. A second round of transfection (LP^2^) was then performed. mCherry-positive cells were again sorted and expanded for two weeks before differentiation. Similarly, a third round of transfection (LP^3^), sorting, expansion, differentiation, and analysis was conducted. B) Experimental design for comparison. FSHD2 iPSCs were subjected to one (LP^1^), two (LP^2^), or three (LP^3^) rounds of transfection with CRISPRoff mRNA and sgRNAs (conditions 3, 6, and 9, respectively). As negative controls, cells were transfected with CRISPRoff mRNA alone for one (Neg^1^), two (Neg^2^), or three (Neg^3^) rounds (conditions 2, 5, and 8, respectively). C) *DUX4-fl*, D) *DUX4* downstream targets, and E) myogenic differentiation markers in cells on day 9 of differentiation. Relative expression was normalized to the average for condition 1 in all samples(n=3). F) DNA methylation analysis of the 5’UTR region of the D4Z4 array by bisulfite sequencing. G-J) Three rounds of transient transfection of CRISPRoff mRNA and sgRNAs efficiently silence *DUX4* in FSHD1 cells. G) Schematic of the transfection, selection, and differentiation workflow for FSHD1 iPSC clones into myocytes. FSHD1 iPSC^tet-MyoD^ clones were transfected by lipofection with CRISPRoff mRNA (mCherry) and sgRNAs for transient expression. One day after seeding, mCherry-positive cells were sorted and expanded for 5 days, followed by second and third rounds of transfection. After three rounds of transfection, mCherry-positive cells were sorted and expanded for two weeks before differentiation for analysis. H) Experimental design for comparison. Three conditions were included: no treatment (NT), FSHD1 iPSCs subjected to three rounds of transfection with CRISPRoff mRNA and sgRNAs (LP^3^), and FSHD1 iPSCs transfected with three rounds of CRISPRoff mRNA alone as negative controls (Neg^3^). I-J) qPCR analysis of *DUX4*-fl and its associated genes, and myogenic differentiation markers in cells on day 9 of differentiation. Relative expression was normalized to the average for condition 1 in all samples(n=3). K) DNA methylation analysis of the 4qA region of the D4Z4 array by bisulfite sequencing. All data are represented as mean ± SEM. *P < 0.05, **P < 0.01, ***P < 0.001, ****P < 0.0001, and ns indicates not significant as determined using one-way ANOVA followed by the Tukey’s multiple comparisons test.

Following successful suppression in FSHD2 cells, we next tested the RNA approach in FSHD1 cells. CRISPRoff mRNA and sgRNAs were transfected into FSHD1 iPSCs three times. mCherry-positive cells were sorted 24 h after the transfection. Cells transfected and sorted three times were then expanded for two weeks before myogenic differentiation (Fig. 5G, H, Supplementary Fig. S5D). *DUX4* and its downstream targets were significantly suppressed after the third induction, confirming *DUX4* silencing in FSHD1 cells (Fig. 5I, J). Since only one 4q allele is hypomethylated due to contraction of the D4Z4 repeat array in FSHD1, the other 4q allele remains hypermethylated. The bisulfite sequencing (BSS) methylation analysis within the D4Z4 repeat units used in FSHD2, cannot clearly distinguish the methylation state of the contracted allele in FSHD1. Instead, as the FSHD1 cells in this study harbored only one 4qA haplotype carrying the contracted D4Z4 array, we performed BSS methylation analysis in the 4qA region rather than in the promoter region within the D4Z4 repeat units [66]. DNA methylation in the 4qA region increased markedly after three rounds of transfection (Fig. 5K, Supplementary Fig. S5E), validating that the hit-and-run RNA approach also effectively suppresses *DUX4* in FSHD1 cells. These results support the potential of RNA delivery to facilitate the clinical translation of hit-and-run epigenetic editing as a therapeutic strategy for all FSHD subtypes.

### CRISPRoff mRNA/sgRNAs-mediated *DUX4*silencing reverses disease-associated gene expression and pathway dysregulation, restoring transcriptome-level alterations

To analyze the genome-wide effects of CRISPRoff-mediated epigenome editing, we performed transcriptomic analysis (RNA-seq) of untreated control (NT), CRISPRoff mRNA alone (Neg^3^), and CRISPRoff mRNA/sgRNAs (LP^3^) conditions. Pairwise comparisons were performed to identify changes genuinely attributable to *DUX4* silencing mediated by epigenome editing. Gene Set Enrichment Analysis (GSEA) based on *DUX4* downstream genes showed a significant overall decrease in LP^3^ cells for genes that have been reported to be upregulated upon *DUX4* expression [64]. Moreover, CRISPRoff alone (Neg^3^) had no significant suppressive effect on *DUX4* downstream genes (Fig. 6B,C). Therefore, the epigenome editing mediated by CRISPRoff mRNA/sgRNAs targeting *DUX4* induced the repression of *DUX4* downstream genes, in line with restoring the repression of *DUX4* downstream genes observed by the treatment with plasmid-based (Fig. 4F) and RNA-based CRISPRoff system (Fig. 5D). In addition, the MA plot for LP^3^ vs. NT showed no significant genome-wide upregulation or downregulation of gene expression, with scattered downregulated genes (Fig. 6D). The top 10 GSEA Hallmark pathways in LP^3^ vs. NT showed negative enrichment of pathways associated with cell proliferation (MYC Targets, E2F Targets, G2M Checkpoint, and Mitotic Spindle), stress response and apoptosis (Hypoxia, Apoptosis, and p53 Pathway), inflammation (TNFA Signaling via NFKB), growth and metabolism (MTORC1 Signaling), and epithelial-mesenchymal transition (EMT) (Fig. 6E). It demonstrated that the upregulation of FSHD-associated pathways involved in RNA processing and cell cycle [11,12], apoptosis [76,77], and inflammation [78] was markedly attenuated upon *DUX4* silencing. In addition, RNA and protein expression analyses in FSHD muscle have shown accumulated *DUX4*-induced oxidative DNA damage, abnormal increase in oxidative stress, and hypoxia perturbation due to mitochondrial dysfunction [78–80].

**Figure 6.**
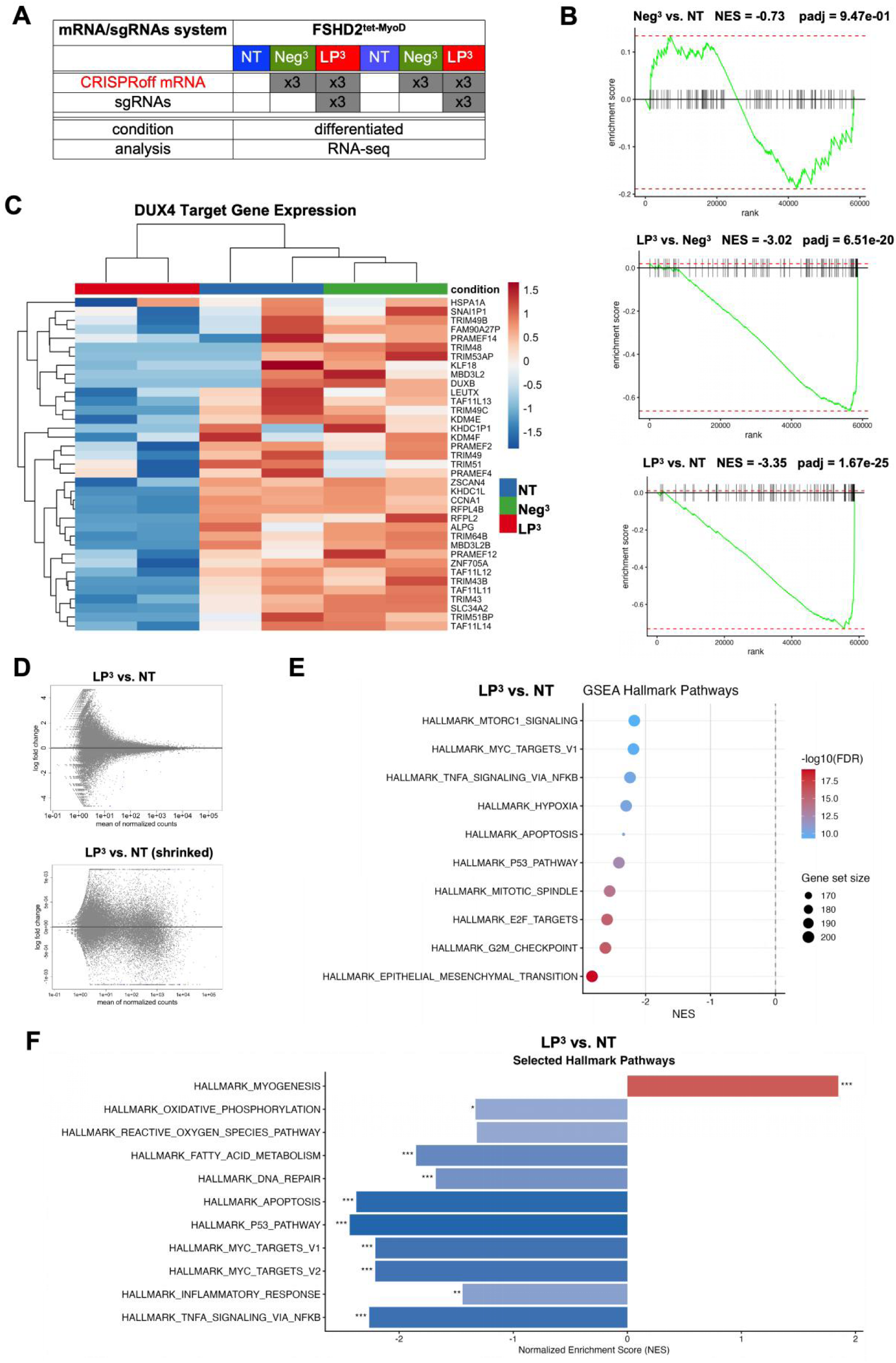
CRISPRoff mRNA-mediated *DUX4* silencing restores *DUX4*-associated gene alteration and FSHD disease-related pathways. See also Supplementary Fig. S6. A) Three conditions were included in this analysis: a) NT: no treatment; b) Neg^3^: CRISPRoff mRNA alone transfected three times; c) LP^3^:CRISPRoff mRNA and gRNAs co-transfected three times. B) Gene Set Enrichment Analysis (GSEA) was performed using the *DUX4* target gene signature. In the CRISPRoff mRNA and sgRNAs co-transfection group (LP^3^), *DUX4* target genes were significantly downregulated compared with both NT (NES=-3.35, padj.=1.67e-25) and Neg^3^ (NES=-3.02, padj.=6.51e-20). In contrast, the CRSIPRoff-alone group (Neg^3^) did not significantly affect *DUX4* target gene expression compared with NT (NES=-0.73, padj.=0.947). NES: normalized enrichment score; padj.: adjusted p-value. C) Heatmap analysis of the leading-edge genes demonstrated that *DUX4* target gene expression was markedly suppressed in the LP^3^ group, whereas NT and Neg^3^ showed no obvious changes. D) MA plots comparing LP^3^ versus NT. The upper panel shows the MA plot using an FDR threshold of 0.05. Grey dots indicate genes without significant changes, whereas blue dots represent significantly downregulated genes. The lower panel shows the MA plot after log-fold-change shrinkage to reduce background noise, revealing that most genes were distributed within ±0.001, close to zero. E) Dot plot visualization of GSEA results for LP^3^ versus NT. The top 10 enriched gene sets ranked by normalized enrichment score (NES) are shown. Each dot represents a gene set, with dot size proportional to the number of enriched genes and color indicating statistical significance (adjusted p-value). F) Bar plots showing hallmark pathways associated with FSHD pathogenesis and *DUX4*-related molecular processes. The selected pathways include myogenesis, oxidative stress and DNA damage, apoptosis, cell proliferation, and inflammation. Results are shown for LP^3^ versus NT.

Consistent with this, GSEA revealed CRISPRoff mRNA/sgRNAs-mediated *DUX4* suppression resulted in negative enrichment of hallmark gene sets for Oxidative Phosphorylation, Reactive Oxygen Species, Fatty Acid Metabolism, and DNA Repair (Fig. 6F). Furthermore, GSEA showed positive enrichment of the hallmark Myogenesis gene set, which suggests that CRISPRoff mRNA/sgRNAs-mediated *DUX4* repression ameliorated the myogenic differentiation defect [13].

Moreover, we compared NT with Neg^3^ to evaluate the changes induced by CRISPRoff alone. The MA plot also showed no obvious genome-wide changes, with only scattered blue dots indicating upregulation (Supplementary Fig. S6A). We observed that the CRISPRoff mRNA treatment was well tolerated by cells as it did not activate apoptosis or impair cell-cycle progression and myogenic differentiation (Supplementary Fig. S6B, E). Interestingly, comparing Neg^3^ with NT, CRISPRoff mRNA alone induced subtle activation of pathways mainly related to oxidative stress and DNA damage, potentially reflecting the reciprocal interaction between *DUX4* expression and oxidative stress [54,77–79] (Supplementary Fig. S6B, E). Although the mechanism of subtle pathway activation mediated by CRISPRoff alone remains unclear, one possible explanation is that hypermethylation may paradoxically promote gene activation by interfering with the binding of transcriptional repressors [81]. Upon targeted *DUX4* silencing, the non-specific transcriptional activation induced by CRISPRoff alone was diminished (Supplementary Fig. S6D, F).

Collectively, these data demonstrate that the CRISPRoff mRNA/sgRNAs-mediated transient intervention strategy not only silences the pathogenic gene and its downstream targets but also restores FSHD-associated pathways, promoting a shift from a diseased, stressful state toward cellular homeostasis.

### CRISPRoff mRNA/sgRNAs method diminished global off-target effect on DNA methylation

We then assessed the specificity of the epigenetic editor by performing Infinium Methylation EPIC array and transcriptome analysis in both the stably integrated dCas9-KRAB plus dCas9-D3A3L (tKD) system and the transient CRISPRoff mRNA/sgRNAs (LP^3^) system. This enabled us to compare off-target effects associated with stable and transient editor expression. The conditions for the two systems are as follows: 1) Stably integrated tKD system (from Supplementary Fig. S3): tKD cells stably expressing dCas9-KRAB(Neo) and dCas9-D3A3L(BFP), and tNN cells stably expressing dCas9-Neg(Neo) and dCas9-Neg(BFP), were subjected to three rounds of transient sgRNA transfection. tNN cells served as the negative control. 2) Transient CRISPRoff mRNA/sgRNAs system (from Fig. 5A-F): LP^3^ cells underwent three rounds of CRISPRoff mRNA and sgRNAs lipofection, while NT cells without treatment served as the negative control (Fig. 7A). PCA analysis showed clear segregation among sample types, confirming the quality of the normalized data (Supplementary Fig. S7A, B). A biased tendency toward upregulation in integrated tKD indicated that 3.26% of probes have potential off-target effects for DNA methylation, consistent with previous reports [46] (Fig. 7B, C, Supplementary Fig. S7C). However, this proportion was markedly reduced to 0.254% in the transient LP^3^ compared with NT (Fig. 7B, C). These results suggest that the transient CRISPRoff mRNA system (LP^3^) substantially reduces the overall hypermethylation tendency induced by the integrative dCas9-effectors (tKD). Then, the differentially hypermethylated probes were annotated to their corresponding genes (Fig. 7D, E, Supplementary Table S5) and integrated with RNA-seq data from differentiated myocytes in the dCas9-effector integrated tEP^3^ (from Fig.2) and transient CRISPRoff mRNA/sgRNAs expressed LP^3^ (from Fig. 5A-F) conditions to assess genome-wide transcriptional off-target effects. In tEP^3^, only two genes (*SCL34A2* and *CCNA1*) were significantly downregulated in the transcriptome (Fig. 7F). *SLC34A2* and *CCNA1* are downstream targets positively correlated with *DUX4* expression in myocytes [11], indicating that the reduced expression of *SLC34A2* and *CCNA1* is a consequence of *DUX4* suppression rather than off-target DNA methylation (Supplementary Fig. S7D). In LP^3^, only *TCEA1* was significantly downregulated (Fig. 7F), potentially because *TCEA1* was predicted as an sgRNA8-related off-target. Taken together, these results demonstrate the specificity of the transient CRISPRoff mRNA/sgRNAs system with minimal off-target methylation and no functional consequences as measured by gene expression.

**Figure 7.**
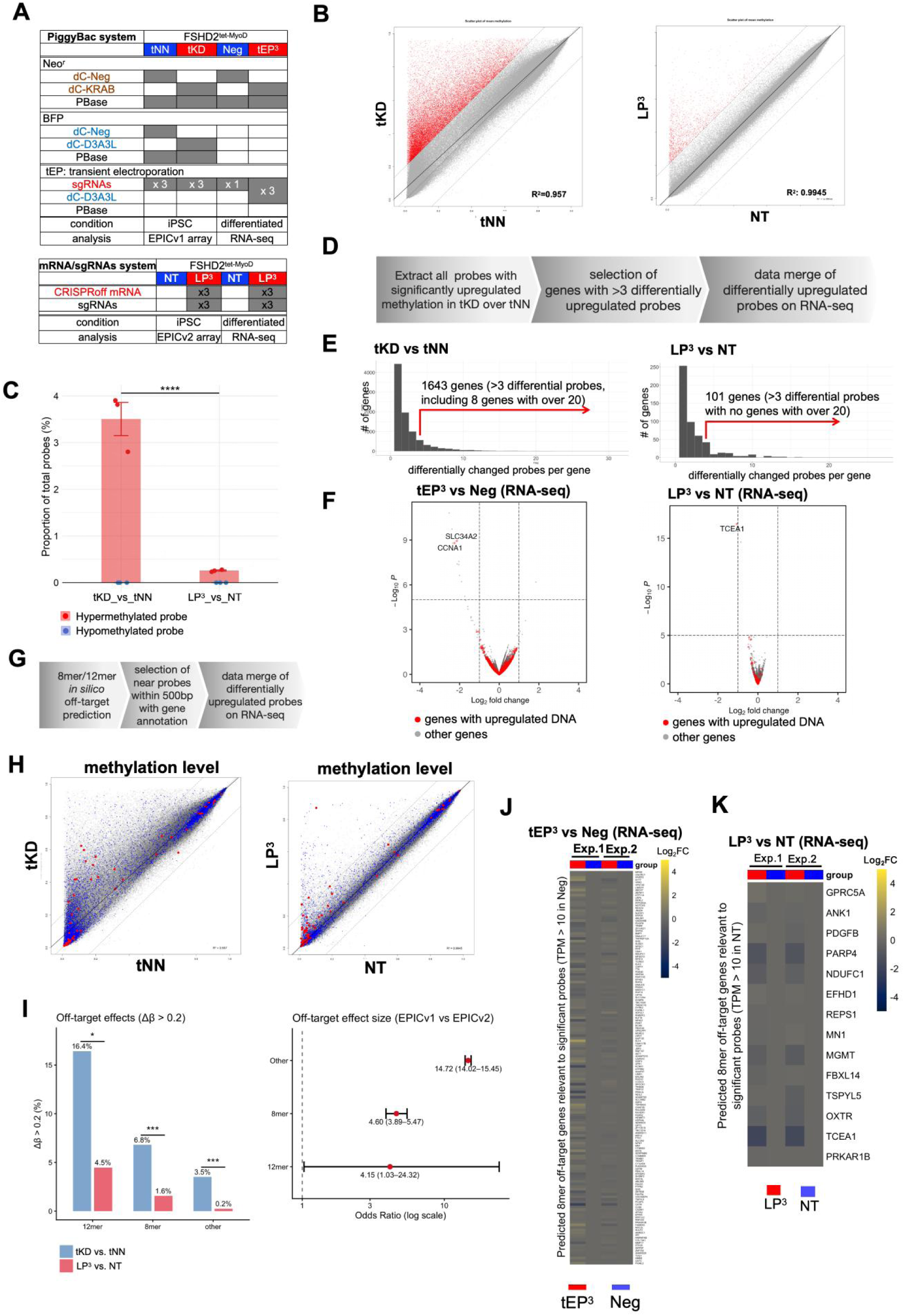
Transient CRISPRoff mRNA/sgRNAs transfection robustly eliminates global off-target effects induced by *piggyBac*-mediated dCas9-effector integration. See also Supplementary Fig. S7. A) Summary of sample conditions used in this analysis. In the *piggyBac* dCas9-effector system, tNN (with dCas9-Neg(Neo) and dCas9-Neg(BFP) constitutively introduced and sgRNAs transiently transfected three times) and tKD (with dCas9-KRAB(Neo) and dCas9-D3A3L(BFP) constitutively introduced and sgRNAs transiently transfected three times) of undifferentiated FSHD2 iPSC^tet-MyoD^ from Supplementary Fig. S3 were used for the Infinium Methylation EPICv1 array. RNA-seq data from the Neg and tEP^3^ of differentiated FSHD2 iPSC^tet-MyoD^ (referred to as conditions 1 and 6 in Fig. 2B) were used. Please note that tNN and tKD correspond to Neg and tEP^3^, respectively, in this analysis. In CRISPRoff mRNA/sgRNAs system, NT (no treatment) and LP^3^ (CRISPRoff mRNA and sgRNAs were transiently transfected three times) in undifferentiated FSHD2 iPSC^tet-MyoD^ (referred conditions 1 and 9 in Fig. 5B) were used for the Infinium Methylation EPICv2 array. The corresponding RNA-seq data from NT and LP^3^ in differentiated FSHD2 iPSC^tet-MyoD^ were obtained. Please note that all Infinium Methylation-associated analysis were based on the 721,347 probes common to EPICv1 and EPICv2. B) The plots of averaged DNA methylation beta values for all common probes in the EPIC array among tKD vs. tNN (n = 2 and 3, respectively) and LP^3^ vs. NT (n=3). when the y-axis compared to the x-axis, significantly upregulated probes with delta-beta>0.2 increased and adj.P-value < 0.01 (up, shown in red) and downregulated probes with delta-beta<-0.2 decreased and adj.P-value < 0.01 (down, shown in blue) are emphasized, with the remaining probes shown as not significant (ns, in grey). Dotted lines show the cutoff with a 0.2 increase or decrease in beta value. C) Mirror-bar Plot on the percentage of hypermethylated and hypomethylated probes in tKD vs. tNN group and LP^3^ vs. NT group from Fig. 7B. Among all common probes, 24,988 of 721,347 (3.26%) showed increased methylation in tKD vs. tNN, compared with 1,835 of 721,347 (0.254%) in LP^3^ vs. NT (Fisher’s Exact Test, OR = 14.06, 95% CI: 13.68–14.45). However, no significant differences were observed when comparing hypomethylated probes. p-value was marked to the comparison (*p-value < 0.05, **p-value < 0.01, ***p-value < 0.001, ****p-value < 0.0001). D) Schematic showing the significant differentially methylated based off-targets. E) Summary of annotated gene numbers with certain numbers of significantly upregulated probes in Fig. 7B. Note that genes with more than three significantly upregulated relevant probes were defined as off-target gene candidates. F) Off-target gene candidates in Fig. 7E were emphasized in red, and the rest in grey, in the corresponding RNA-seq data. G) Schematic showing the in silico off-target prediction-based evaluation. See also supplementary Fig. 7E for a detailed workflow of the in silico off-target prediction. H) Probes proximal to each predicted 8-mer (in blue) and 12-mer (in red) off-target binding site for all sgRNAs were emphasized in all dots (same as Fig. 7B) of the averaged EPIC array beta value in tKD vs tNN group and LP^3^ vs. NT group. I) Bar Plot showing the percentage of delta-beta>0.2 in tKD vs. tNN (blue column) and the LP^3^ vs. NT (red column). Among the 8-mer off-target genes, 763 of 11,214 (6.8%) were hypermethylated in tKD, compared with 175 of 11,215 (1.6%) in LP^3^ (Fisher’s Exact Test, OR = 4.60, 95% CI: 3.89–5.47). Similarly, among the 12-mer off-target genes, 11 of 67 (16.4%) were hypermethylated in tKD, whereas only 3 of 67 (4.5%) were hypermethylated in LP^3^ (Fisher’s Exact Test, OR = 4.15, 95% CI: 1.03–24.32). p-value was marked to each comparison (*p-value < 0.05, **p-value < 0.01, ***p-value < 0.001). The odds ratio and 95% confidence interval from Fisher’s Exact Test were depicted in a forest plot. J) The annotated genes with at least one significantly upregulated probe (> 0.2) from the off-target proximal probes in Fig. 7H were merged using RNA-seq data, with relative changes shown using a heatmap for each experimental set. The gene list for Heatmap was shown in Supplementary Table S7.

Next, we further analyzed the in silico off-target effects associated with sgRNAs. First, we used CRISPR direct [69] to predict sgRNA off-target sites. Then, we mapped the corresponding off-target sites to the genome, screened for probes near off-target sites, and added annotation information to identify off-target genes (Supplementary Fig. S7E-H, Supplementary Table S6). Hypermethylation of 12-mer off-target genes in the transient CRISPRoff mRNA/sgRNAs (LP^3^) system was significantly reduced compared with the stably integrated tKD system. In tKD, 6.8% of 8-mer off-target genes showed hypermethylation, while this proportion decreased to 1.6% in LP^3^ (Fig. 7H, I). In addition, 8-mer off-target genes with differentially hypermethylated probes were integrated with RNA-seq data from relevant differentiated myocytes to see identify gene expression changes. Regardless of methodology, the influence on gene expression of off-target genes was minimal (Fig. 7J, K, Supplementary Table S7). Collectively, these data suggest that transient RNA delivery of epigenome-editing components substantially reduces hypermethylation at predicted off-target sites while having limited impact on the transcriptome.

### CRISPRoff-mediated hit-and-run successfully suppressed *DUX4*in iPSC-derived muscle stem cells

We confirmed the suppressive effects of the hit-and-run strategy on *DUX4* expression in iPSC models. However, this strategy needs to be evaluated in muscle progenitor cells and throughout their differentiation into mature myotubes to explore its therapeutic potential for clinical application. Here, we generated iPSC-derived muscle stem cells (iMuSCs) from FSHD2 iPSCs and its isogenic control—in which the disease-causing *SMCHD1* mutation was corrected—using a stepwise differentiation protocol established by our lab [70], allowing us to evaluate whether the hit-and-run strategy could efficiently suppress *DUX4* in muscle stem cells. To investigate the expression profiles during iMuSCs differentiation, we characterized expression changes in *DUX4* and its downstream genes at day 0 and every 2 days from day 4 to day 18 (Supplementary Fig. S8A). Compared with the isogenic control, FSHD2 iMuSCs exhibited distinct expression dynamics of *DUX4* and its downstream genes. The expression levels of *DUX4* and its downstream genes increased rapidly during the first week of differentiation, followed by a slower increase during the second week. By approximately day 16, the expression of *DUX4* and its downstream genes appeared to reach a plateau (Supplementary Fig. S8B). These results validate our iMuSCs model by demonstrating that *DUX4* and downstream genes are not expressed during the muscle stem cell proliferation stage but are activated upon myogenic differentiation in FSHD2 iMuSCs, while remaining undetectable in the isogenic control. To determine whether the CRISPRoff mRNA/sgRNAs-mediated hit-and-run silencing strategy results in sustained *DUX4* suppression, FSHD2 iMuSCs were subjected to three consecutive rounds of CRISPRoff mRNA/sgRNAs lipofection, followed by one week of proliferation and subsequently one week of myogenic differentiation. The expression of *DUX4* and its downstream genes was then analyzed (Supplementary Fig. S8C). The expression levels of *DUX4* and its downstream genes were reduced in the CRISPRoff mRNA/sgRNAs lipofection-treated (LP^3^) group compared with the untreated group (NT), without affecting myogenic marker gene expression (Supplementary Fig. S8E-G). These findings confirm that muscle stem cells can also be effectively targeted using the CRISPRoff-mediated hit-and-run silencing strategy.

## DISCUSSION

In this study, we revealed that hit-and-run silencing targeting the D4Z4 array can suppress patient-specific endogenous *DUX4* expression beyond the distinct pathogenic genotypes of FSHD1 and FSHD2. Importantly, to the best of our knowledge, this is the first report demonstrating that transient RNA delivery of an epigenetic editor enables sustained gene silencing targeting the repeat array by restoring congenital DNA hypomethylation of the repeat array to a hypermethylated state comparable to that of unaffected cells, ultimately restoring dysregulated disease-associated pathways toward a normal state. While constitutive expression of single dCas9-KRAB, dCas9-D3A, or dCas9-D3A3L could increase DNA methylation to a limited range on the D4Z4 array, transient co-delivery of combined KRAB and D3A3L produced a stronger methylation effect that spreads into regions distal to sgRNA target sites. These findings suggest a synergistic interaction between the two effectors that cannot be achieved by either effector alone, consistent with previous studies targeting a single genomic locus [50,52]. Unlike previous reports in which a single transient transfection was sufficient to induce hit-and-run silencing at a single genomic locus, our approach required multiple rounds of hit-and-run to achieve robust *DUX4* suppression. This may be because multiple D4Z4 repeat units were targeted simultaneously throughout the entire genome, effectively diluting the available enzymatic activity and limiting the extent of methylation at each repeat unit from a single treatment. This study provides evidence that hit-and-run is not limited to the modulation of a single locus but can also achieve sustainable editing across multiple repetitive regions. The D4Z4 array in unaffected individuals is epigenetically characterized not only by DNA hypermethylation but also by accumulation of the heterochromatin marker H3K9 trimethylation or HP1 gamma [82], and forms distinct genome structures outside the repeats [83–87], which are reduced and abnormally rearranged, respectively, in FSHD. As our epigenetic silencing strategy achieved D4Z4 array hypermethylation comparable to unaffected levels, it would be interesting to further investigate whether the strategy could reconstruct unaffected-like heterochromatin structures in muscle cells directly through DNA hypermethylation, whose role in pathology remains controversial [88].

Several potential methods for FSHD therapy have been reported, which primarily focus on suppressing *DUX4* transcription or translation. Chemical compounds targeting RNA interference or signal transduction can effectively suppress *DUX4* expression [20–25]; however, their therapeutic application may be limited by poor molecular stability, reduced specificity, the need for repeated long-term administration, and potential off-target effects arising from systemic modulation of multifunctional signaling pathways. By contrast, a hit- and-run silencing strategy may overcome some of these limitations by enabling sustainable target gene suppression after a single or limited intervention, thereby reducing the need for repeated administration, hospital visits, and treatment-related financial burdens and ultimately improving FSHD patients’ quality of life. Genome editing should be further tested to better understand permanent transcriptional repression, but it remains technically challenging to edit the D4Z4 array. While editing exon3(PAS) outside the D4Z4 repeat unit is technically promising for treatment, two groups have unfortunately observed an unexpected *DUX4* mRNA switch phenomenon, whose underlying mechanism remains unclear [26,27].

Given this unexpected transcript switch and concerns about potential on- and off-target effects that may lead to unintended mutations, the epigenetic silencing strategy described in this study may provide a more effective, safer, and more controllable approach to gene silencing without directly altering the genomic sequence. Moreover, clinical studies of large FSHD patient cohorts demonstrated that 4qA-specific distal D4Z4 hypomethylation is associated with disease severity and progression [31,32], supporting our concept that restoring D4Z4 methylation may help prevent FSHD disease progression. This strategy’s potential therapeutic benefit may be further enhanced by initiating treatment at the onset of early symptoms. Early intervention may help slow disease progression and preserve muscle function before substantial muscle damage occurs. Our current understanding of the mechanisms underlying *DUX4* activation, particularly in FSHD muscle cells, is limited.

Several signaling pathways have been revealed in a context-dependent manner [55,89–93]. In this sense, reconstructing chromatin structure in patient nuclei using our approach may be a more inclusive way to render the *DUX4* locus insensitive to any environmental stimulation. Since DUX4 is strongly activated during differentiation and muscle regeneration [12,94], our approach may be effective for patients experiencing excessive exhaustion or muscular damage even when they cannot access chemical compound drugs.

EPIC array analysis was performed to evaluate off-target DNA methylation changes associated with constitutive vector-based expression and transient RNA delivery of epigenetic editing components. Transient transfection markedly reduced off-target DNA methylation effects compared with constitutive transfection, suggesting that limiting editor exposure temporally effectively decreases “floating” enzymatic off-target effects. Integrated analysis of EPIC array and RNA-seq data revealed that hit-and-run DNA methylation targeting had only minimal effects on the transcriptome at non-targeted loci. In the transient RNA system, we observed mild downregulation of *TCEA1* (Transcription Elongation Factor A1), a predicted off-target gene of sgRNA8. *TCEA1* is a human gene encoding the transcription elongation factor that plays a critical role in regulating mRNA synthesis. The downregulation was also potentially because the disruption of mRNA synthesis regulation [8,9] was eliminated following *DUX4* silencing (Fig.6E, F). Moreover, although we simultaneously applied four validated sgRNAs targeting D4Z4 to establish proof of concept, sgRNAs that maximize potency to robustly suppress *DUX4* while minimizing off-target effects should be prioritized for clinical applications. This approach may also help reduce sgRNA-guided binding-related off-target effects. Furthermore, intergenic regulatory regions have recently been shown to have pathogenic potential due to disrupted fine-tuning of gene expression, especially in humans [95,96]. In this sense, in the context of future clinical use, higher-coverage methods such as bisulfite high-throughput sequencing-based or long-read sequencing-based off-target analysis can be applied to strictly evaluate off-target risks in a sustainable manner to allow for future re-analysis for *de novo* identified naive sites.

We have shown that hit-and-run silencing functioned in FSHD patient-derived cells in a proliferative pluripotent state, and the effect was toleranted during cell divisions for at least two weeks (maximally 10 weeks in our single trial, data not shown) and muscle differentiation, suggesting that epigenetically robust and inheritable chromatin structures were maintained across cell types. For the clinical application, it may be necessary to confirm whether hit-and-run silencing can be achieved with long-term benefits in various myogenic cells residing in the skeletal muscle tissues, such as quiescent satellite cells, proliferative myoblasts, and terminally differentiated myofibers. Using an iMuSC model generated from FSHD2 patient-derived iPSCs that recapitulated the *DUX4* expression profile observed in primary muscle cells, we further validated hit-and-run silencing to target the D4Z4 array for *DUX4* suppression, demonstrating this approach in satellite-like cells. Given the spike-like activation of *DUX4* in a sporadic manner during differentiation in FSHD muscle cells and regeneration in an FSHD mouse model containing an FSHD patient-derived contracted D4Z4 array [12,94], high transfection efficiency and sustainable effects in satellite cells will be required to suppress disease progression in FSHD muscles. Although AAV vectors are a promising delivery method into muscle tissues due to their high efficiency [43], they have limited packaging capacity. Moreover, recent reports have raised potential risks in clinical use in terms of lethal liver dysfunction and immunogenicity [42,97]. Considering that one of the benefits of hit-and-run silencing is that there is no need for dCas9/sgRNAs components to remain in the nuclei to produce long-term effects, it has high potential in clinical application when combined with less toxic *in vivo* transfection methods [98,99], as it allows for the removal of any artificial components from patient bodies after induction. In this sense, customized local applications targeting critical muscles based on each patient’s pathophysiology rather than a systemic approach may be sufficient [100] and are helpful to minimizing side effects in other tissues. In addition, because the hit-and-run epigenetic approach does not manipulate DNA sequences, it likely leads to fewer ethical concerns than with direct genetic approaches, as it cannot produce artificial sequences through unnatural processes [28,101].

In summary, we provide evidence that the hit-and-run silencing strategy targeting the D4Z4 repeat array can effectively reduce *DUX4* expression in FSHD patient-derived cells *in vitro*, with minimal detectable off-target effects. This approach may offer a promising strategy to overcome many challenges associated with conventional therapeutic modalities and achieve more targeted, sustainable treatment for individuals with FSHD. Further optimization in relevant mouse models combining suitable delivery methods [73–75,102] will be essential to establish proof of concept, evaluate the long-term efficacy and safety of this approach, and ultimately assess its potential for clinical translation.

## ACKNOWLEDGEMENTS

We are grateful to the donors who provided their cells for the establishment of iPSCs used in this study. We thank Mr. Masaki Nomura, Mrs. Hiromi Dohi and Mrs. Maki Hira at the faculty of CiRA Foundation for performing the DNA methylation EPIC array protocol. We thank Dr. Akitsu Hotta (CiRA, Kyoto), Dr. Stanly Qi (Stanford University, California), Dr. Peter L. Jones (University of Nevada, Nevada) and Dr. Jun Otomo (CiRA, Kyoto) for critical comments on our study; Ms. Kazusa Okita (CiRA, Kyoto) for technical assistance with the RNA-seq experiment; Dr. Kanae Mitsunaga (CiRA, Kyoto) for technical support of FACS; Dr. Kelvin Hui (CiRA, Kyoto) for English proofreading; and Dr. Chikako Okubo (CiRA, Kyoto) for technical support of NGS data analysis. “GRAPHICAL ABSTRACT” created in BioRender.

He, J. (https://BioRender.com/vks7wjg) is licensed under CC BY 4.0. ChatGPT (OpenAI, GPT-6) was used to assist with editing and improving R code and to refine the wording of the manuscript. The authors directed the use of the tool, reviewed and edited ChatGPT-generated content, and verified the accuracy and integrity of the manuscript. We appreciate the English language editing services provided by Editage (www.editage.com). We appreciate all members of the Sakurai Laboratory for their kind support. We especially appreciate our families and the Japan FSHD patient community for generous and unwavering support.

## Author Contributions

J.H. (Conceptualization [Equal], Data curation [Lead], Formal analysis [Lead], Funding acquisition [Lead], Investigation [Lead], Methodology [Lead], Software [Lead], Validation [Lead], Visualization [Lead], Writing - original draft [Lead], Writing - review & editing [Lead]), M.S.-H. (Conceptualization [Lead], Data curation [Lead], Formal analysis [Lead], Funding acquisition [Lead], Investigation [Equal], Methodology [Lead], Project administration [Equal], Resources [Equal], Software [Lead], Supervision [Lead], Validation [Equal], Visualization [Lead], Writing - original draft [Equal], Writing - review & editing [Lead]), T.J. (Investigation [Equal], Methodology [Supporting], Resources [Supporting], Validation [Supporting], Writing - review & editing [Supporting]), M.A. (Investigation [Equal], Methodology [Supporting], Validation [Equal], Writing - review & editing [Supporting]), T.K. (Data curation [Supporting], Investigation [Equal], Validation [Supporting], Writing - review & editing [Supporting]), A.M. (Data curation [Supporting], Investigation [Equal], Methodology [Supporting], Resources [Supporting], Validation [Supporting], Writing - review & editing [Supporting]), C.M. (Data curation [Supporting], Investigation [Equal], Methodology [Supporting], Writing - review & editing [Supporting]), S.I.(Data curation [Supporting], Investigation [Equal], Methodology [Supporting], Resources [Supporting], Writing - review & editing [Supporting]), S.S.(Data curation [Supporting], Formal analysis [Supporting], Methodology [Supporting], Writing - review & editing [Supporting]), T.Y.(Formal analysis [Supporting], Investigation [Supporting], Methodology [Supporting], Resources [Supporting], Writing - review & editing [Supporting]), H.S. (Conceptualization [Lead], Data curation [Lead], Funding acquisition [Lead], Investigation [Supporting], Methodology [Lead], Project administration [Lead], Resources [Lead], Supervision [Lead], Validation [Supporting], Writing - original draft [Supporting], Writing - review & editing [Lead]).

## SUPPLEMENTARY DATA

Supplementary Data are available at NAR online.

## DATA AVAILABILITY

All RNA-seq data and EPIC array data used in this study are available in the NCBI GEO database (accession number: GSE201185, GSE346195, GSE346307)

