## SUPPLEMENTARY MATERIAL for "Hit-and-run silencing of endogenous *DUX4* by targeting DNA hypomethylation on D4Z4 repeat array in facioscapulohumeral muscular dystrophy"

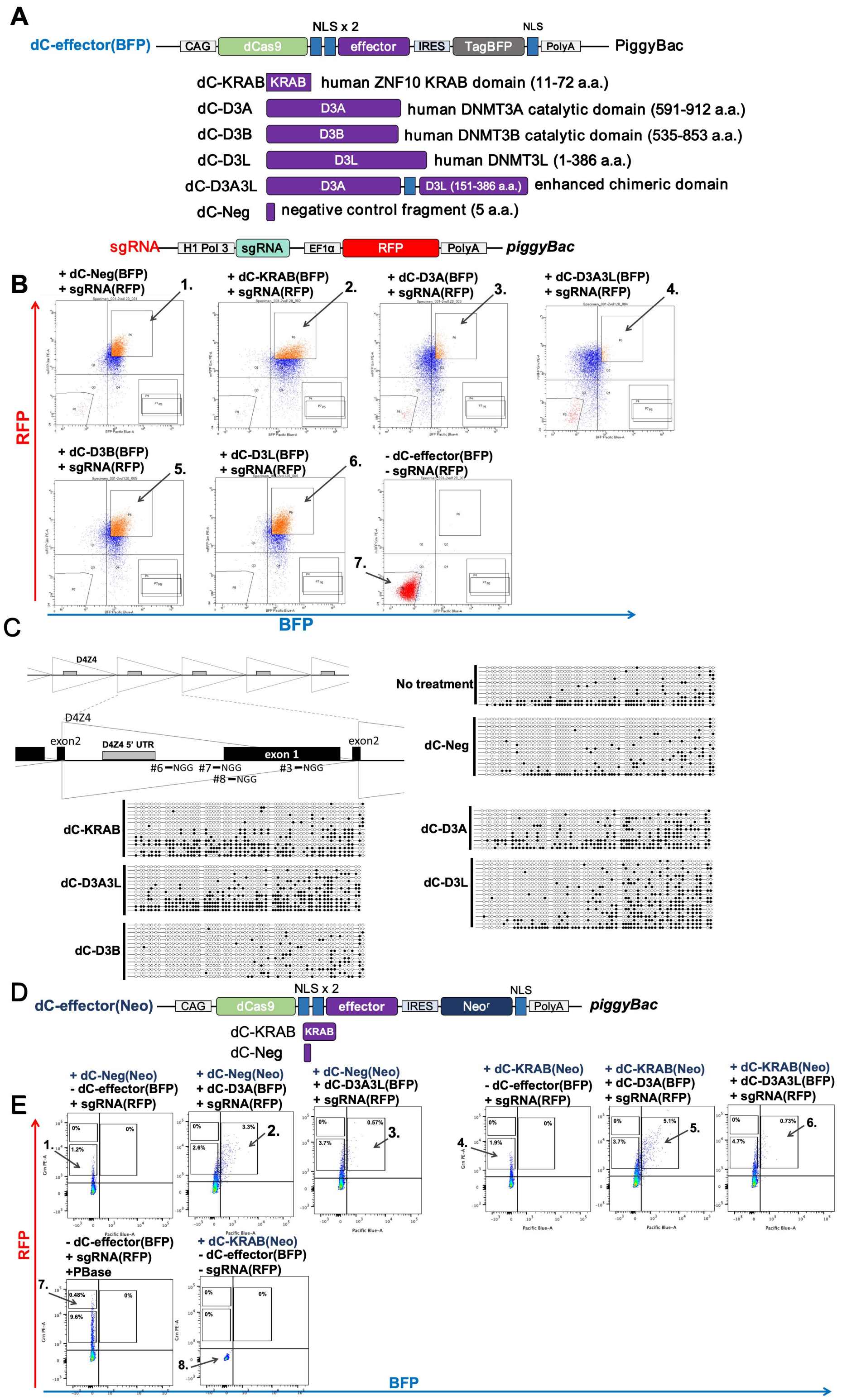


Supplementary Fig. S1. related to Fig.1

A) Construct designs of the *piggyBac* vectors used in Fig.1A. dCas9-effectors are driven by a CAG promoter and followed by internal ribosome entry site (IRES) linked to a blue fluorescent protein (TagBFP) reporter. The sgRNA vectors contain an red fluorescent protein (RFP) reporter. B) Representative images of sorting gates one week electroporation with the dCas9-effector (BFP) and sgRNA (RFP) vectors under constitutive-expression conditions, corresponding to Fig. 1A. Cells were sorted from the square gates indicated by arrows under each condition. C) D4Z4 CpG methylation status of each condition, determined by bisulfite sequencing (BSS), to confirm the enzymatic activity in Fig. 1E. The grey rectangle indicates the region corresponding to the 5' UTR of *DUX4* analyzed by BSS. D) Construct designs of the dCas9-effector vectors, in which a neomycin resistance marker for drug selection is expressed under the control of an IRES. E) Representative images of sorting gates two days after transient electroporation with dCas9-effector(BFP) and sgRNA(RFP) vectors, corresponding to Fig. 1F. The square gates indicated by arrows show the populations sorted under each condition for further expansion. Note that condition 1–6 showed almost no cells with very high RFP signals, unlike condition 7, which underwent constitutive transfection.

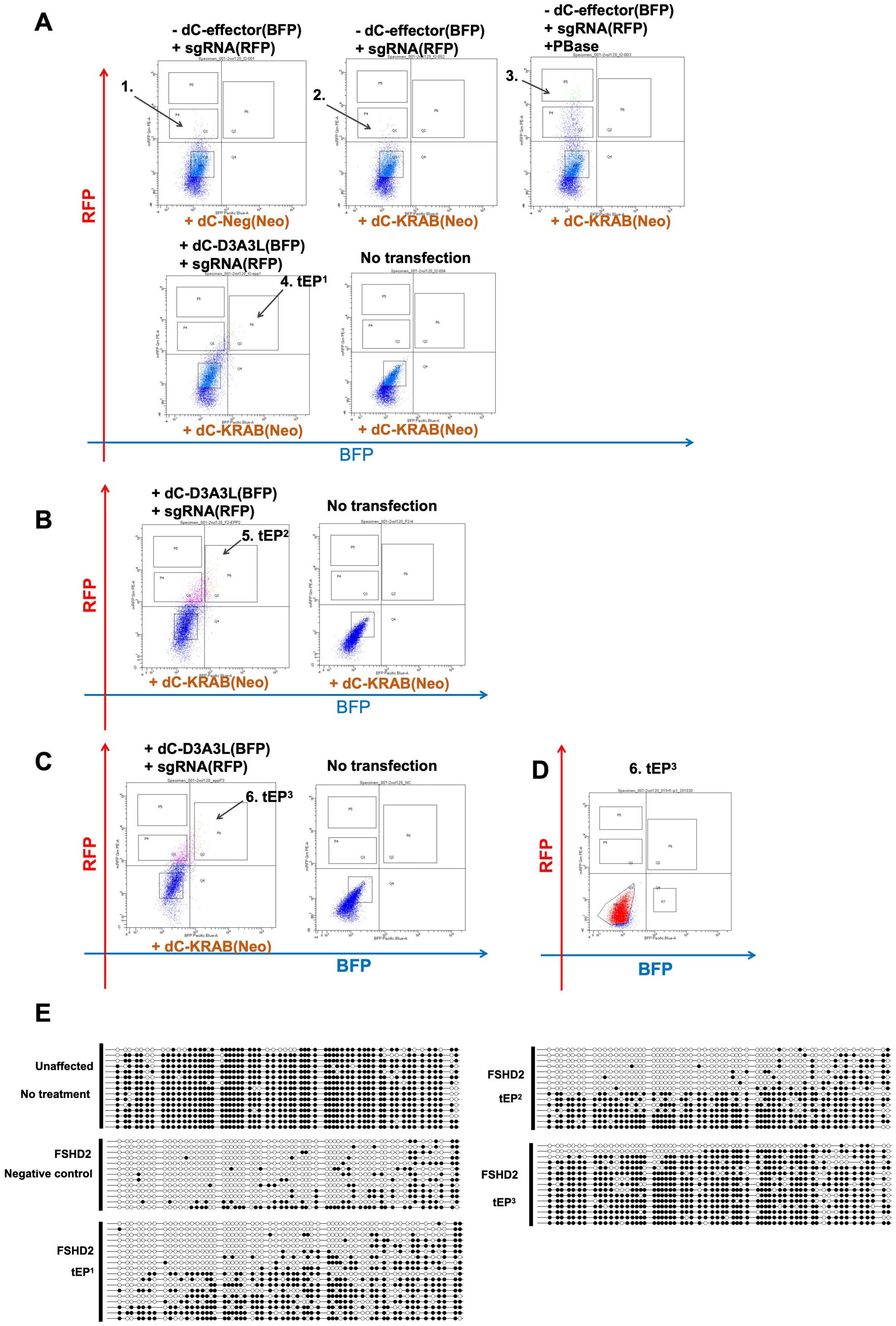


Supplementary Fig. S2. Related to Fig.2

A–D) Representative images of sorting gates corresponding to Fig. 2A. The square gates indicated by arrows show the cell populations sorted under each condition. A) Two days after the first round of transfection. B) Two days after the second round of transfection. C) Two days after the third round of transfection. D) Seven days after the third transfection, when dCas9-D3A3L (BFP) and sgRNAs (RFP) signals were no longer detectable. E) D4Z4 CpG methylation status of individual clones, determined by bisulfite sequencing (BSS), corresponding to the analysis shown in Fig. 2I. For the FSHD2 negative control, cells were treated with tEP[1], and cells negative for transfection markers were sorted.

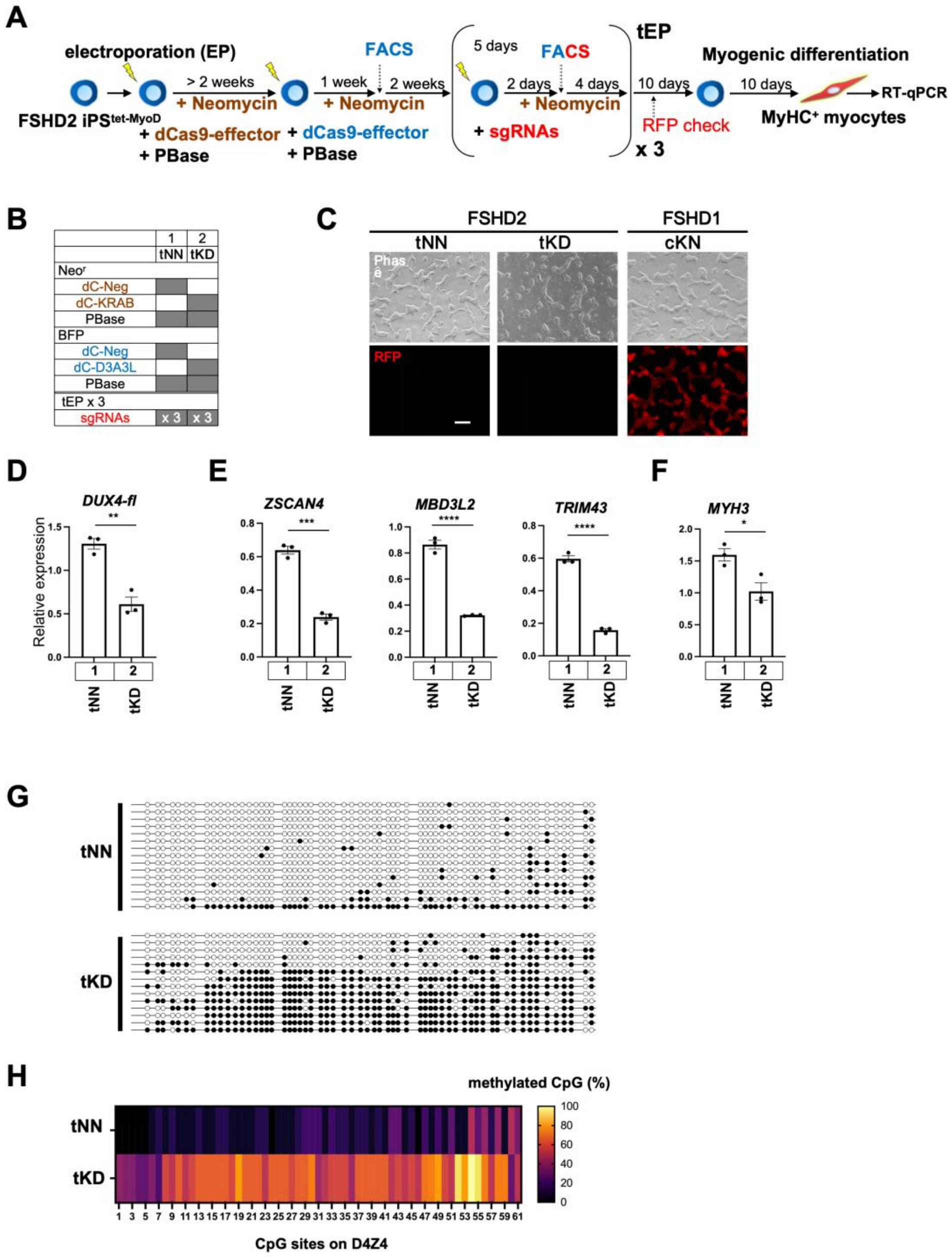


Supplementary Fig. S3. related to Fig.3

A-H) Validation of the modified transfection strategy with FSHD2-iPSC model.

A) Scheme of the transfection, selection and differentiation workflow for FSHD2-iPSC clones into myocytes. FSHD2 iPSC$^{tet\text{-}MyoD}$ clones were first transfected with the dCas9-effector (Neo) vector and PBase to establish constitutive expression. Subsequently, the dCas9-effector (BFP) was integrated using PBase, and BFP-positive cells were sorted. After more than two passages to allow the removal of residual PBase following electroporation, cells were transfected with sgRNAs without

PBase to achieve transient expression. Two days after transfection, BFP- and RFP-positive cells were collected by cell sorting and expanded for four days. This transfection-and-sorting procedure was performed for three rounds, followed by two weeks of expansion and subsequent differentiation for analysis. Note that the dCas9-effector is constitutively expressed, whereas its targeted activity is induced only upon sgRNA delivery. B) Experimental design for comparison. C) Representative phase-contrast and RFP fluorescence image obtained one week after the third round of transfection. cK cells derived from FSHD1-iPSCs were used as a positive control. D-F) RT-qPCR analysis of D) *DUX4-fl*, E) *DUX4* downstream target genes, and F) a myogenic differentiation marker in cells on day 10 of differentiation (n=3). All RT-qPCR data are represented as mean±SEM. *P < 0.05, **P < 0.01, ***P < 0.001, ****P < 0.0001, and ns indicates not significant, as determined using unpaired Student’s t-test. G-H) DNA methylation analysis by bisulfite sequencing (BSS) to confirm enzymatic activity. G) D4Z4 CpG methylation status of individual clones. H) Heatmap showing the average CpG methylation status.

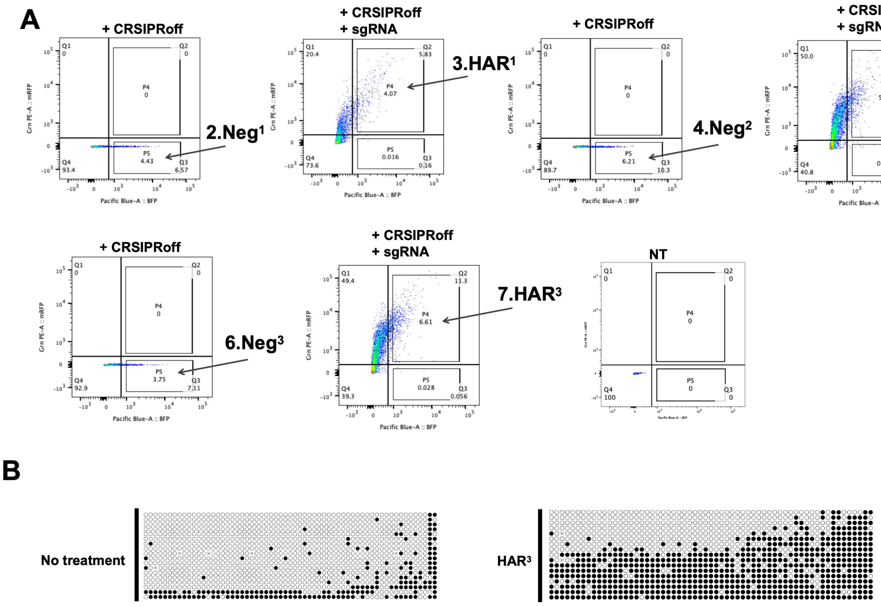


Supplementary Fig. S4. related to Fig.4

A) Representative images of sorting gates corresponding to Fig. 4A. Square gates indicated by arrows represent the sorting gates for each condition at day 2 post-transfection, from which cells were collected for further expansion. B) D4Z4 CpG methylation status of individual clones, determined by BSS, confirming enzymatic activity shown in Fig. 4H.

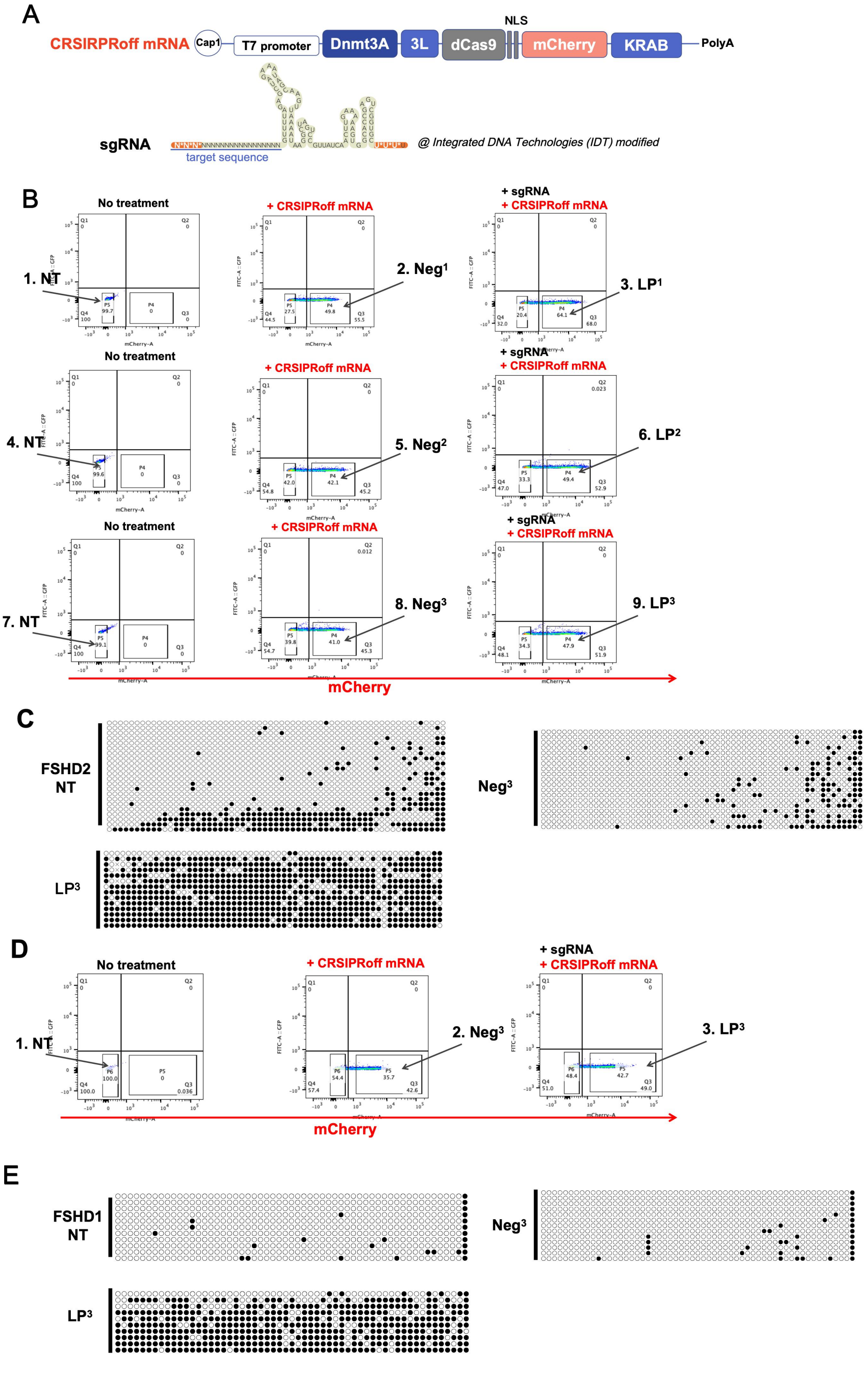


Supplementary Fig. S5. related to Fig.5

A) Construct designs for the CRISPRoff mRNA and sgRNA used in Fig.5A. The N¹-methylpseudouridine (m¹Ψ)-modified CRISPRoff mRNA encodes DNMT3A-DNMT3L, dCas9 and KRAB elements, with a red fluorescent protein (mCherry) reporter. The sgRNA construct designs show bold white letters indicating the 2’-O-methyl (2’-OMe)-modified nucleotides, and asterisks representing the phosphorothioate (PS) linkage. These chemical modifications were introduced to improve RNA stability and nuclease resistance. The target sequences of the sgRNAs are identical to those used in the vector system. B) Representative images of sorting gates at 24 h after each round of transfection in the FSHD2-iPSC model. The square gates indicated by arrows mark the cell populations sorted for subsequent expansion. C) D4Z4 5’UTR CpG methylation status of individual clones, determined by BSS, confirming enzymatic activity (Fig. 5F). D) Representative images of sorting gates for FSHD1 cells at 24 h after the third round of transfection. The same gating strategy was applied across all three rounds. The square gates indicated by arrows mark the cell populations sorted for subsequent expansion. E) D4Z4 4qA CpG methylation status of individual clones, determined by BSS, confirming enzymatic activity (Fig. 5K).

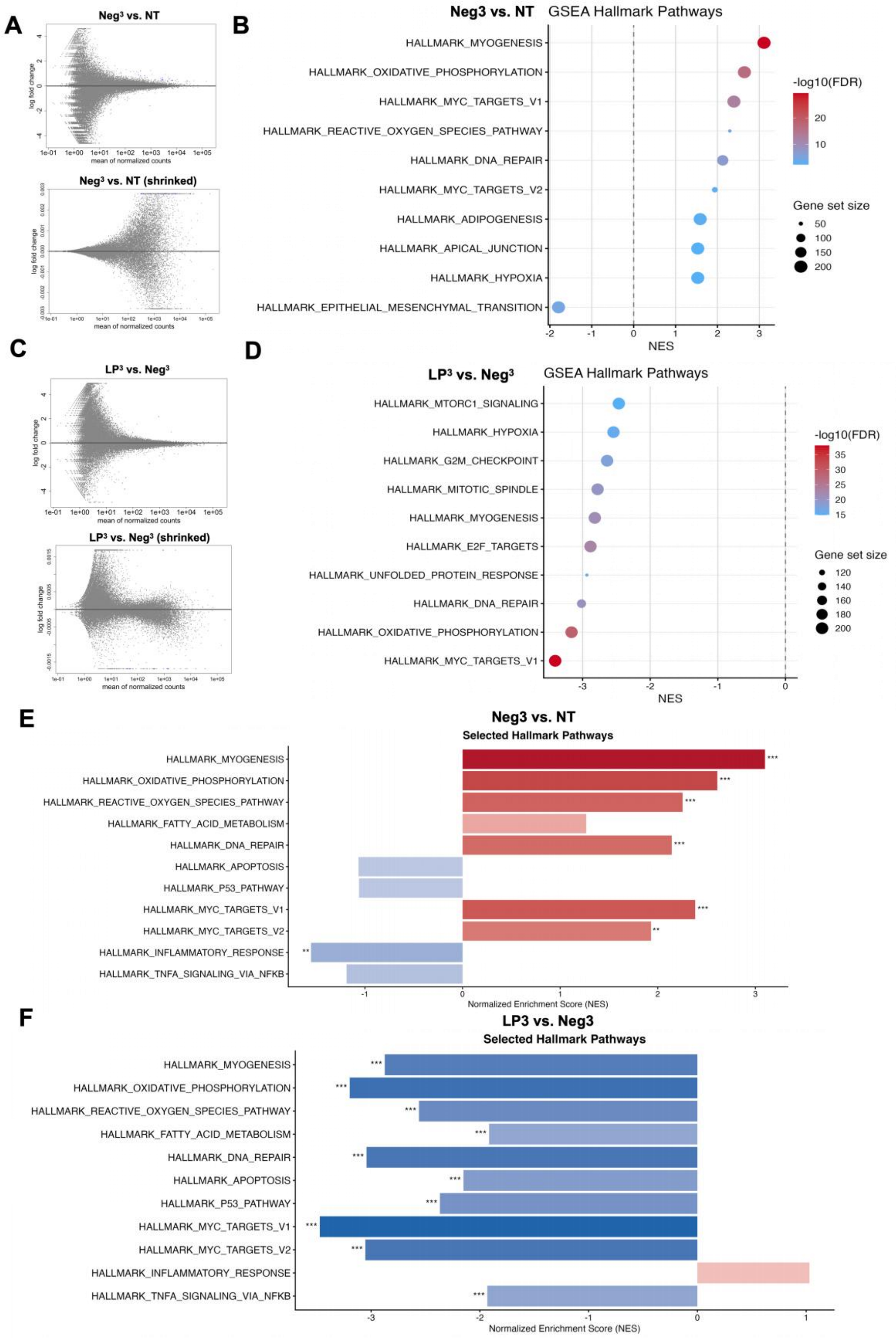


Supplementary Fig. S6. related to Fig. 6.

A) MA plots comparing Neg$^3$ versus NT. The upper panel shows the MA plot using an FDR shreshold of 0.05. A few scattered blue dots represent significantly upregulated genes. The lower panel shows the MA plot after log-fold-change shrinkage, demonstrating that most genes were distributed close to zero (within±0.003). B) Dot plot visualization of GSEA results for Neg$^3$ versus NT. The top 10 enriched gene sets ranked by normalized enrichment score (NES) are shown. Each dot represents a gene set, with dot size indicating the number of enriched genes and color representing the adjusted p-value. C) MA plots comparing LP$^3$ versus Neg$^3$. The upper panel shows the MA plot using an FDR threshold of 0.05. A few scattered blue dots represent significantly downregulated genes. The lower panel shows the MA plot after log-fold-change shrinkage, demonstrating that most genes were distributed within±0.0015. D) Dot plot visualization of GSEA results for LP$^3$ versus Neg$^3$. The top 10 enriched gene sets ranked by normalized enrichment score (NES) are shown. Each dot represents an enriched gene set, with dot size indicating the number of enriched genes and color intensity reflecting the adjusted-p value. E-F) Bar plots showing hallmark pathways associated with FSHD pathogenesis and *DUX4*-related molecular processes. The selected pathways include myogenesis, oxidative stress and DNA damage, apoptosis, cell proliferation, and inflammation.Results are shown for E) Neg$^3$ versus NT and F) LP$^3$ versus Neg$^3$.

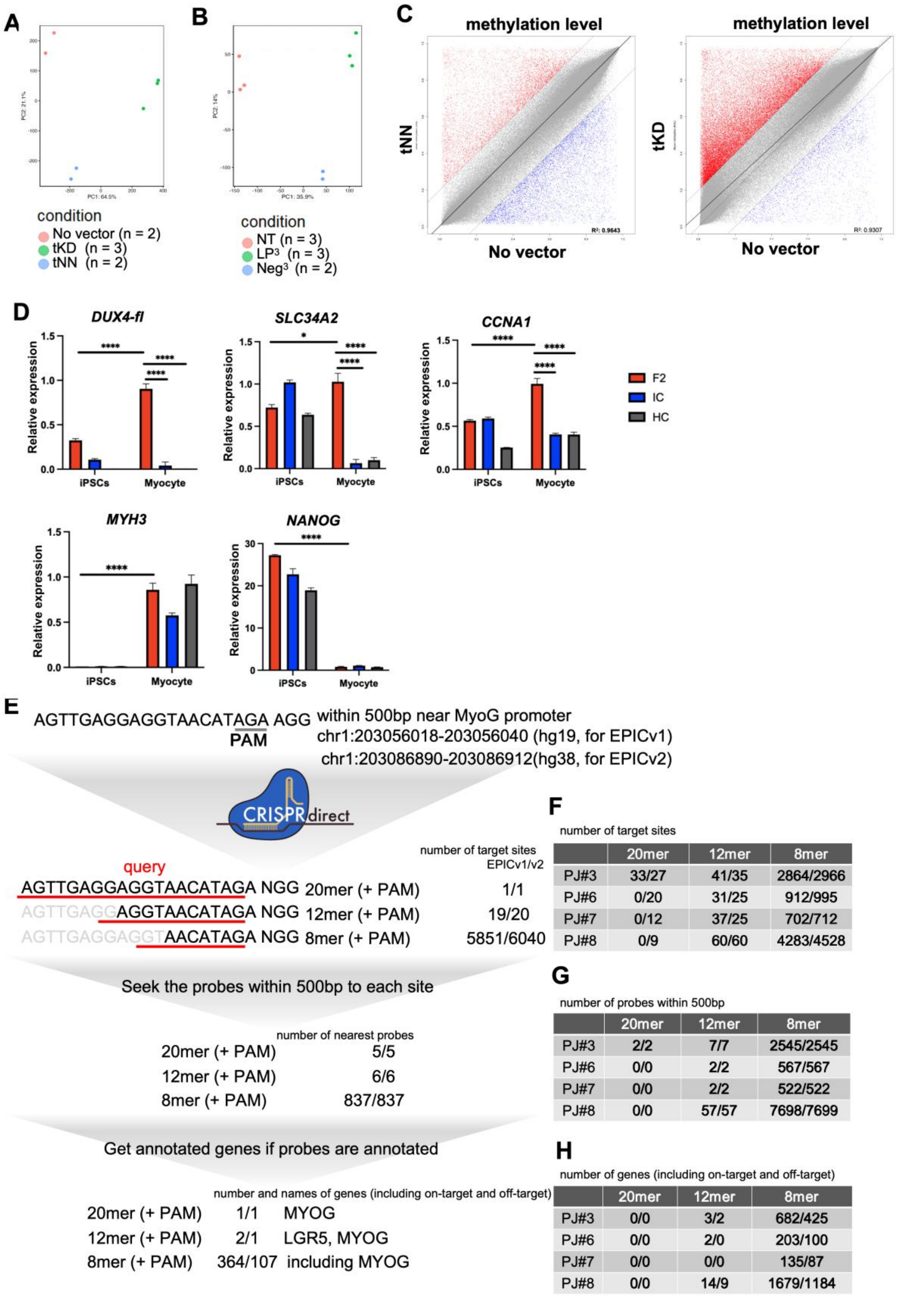


F — number of target sites

| | 20mer | 12mer | 8mer |
|---|---|---|---|
| PJ#3 | 33/27 | 41/35 | 2864/2966 |
| PJ#6 | 0/20 | 31/25 | 912/995 |
| PJ#7 | 0/12 | 37/25 | 702/712 |
| PJ#8 | 0/9 | 60/60 | 4283/4528 |

G — number of probes within 500bp

| | 20mer | 12mer | 8mer |
|---|---|---|---|
| PJ#3 | 2/2 | 7/7 | 2545/2545 |
| PJ#6 | 0/0 | 2/2 | 567/567 |
| PJ#7 | 0/0 | 2/2 | 522/522 |
| PJ#8 | 0/0 | 57/57 | 7698/7699 |

H — number of genes (including on-target and off-target)

| | 20mer | 12mer | 8mer |
|---|---|---|---|
| PJ#3 | 0/0 | 3/2 | 682/425 |
| PJ#6 | 0/0 | 2/0 | 203/100 |
| PJ#7 | 0/0 | 0/0 | 135/87 |
| PJ#8 | 0/0 | 14/9 | 1679/1184 |

Supplementary Fig.S7. related to Fig.7

A-C) Validation of data quality of the Infinium MethylationEPIC array. As for *piggyBac* system, FSHD2 iPSCs from a different donor without any dCas9 or sgRNA vectors (No vector) was also included as a control in analysis. A-B) Principle component analysis (PCA) of individual samples based on normalized beta values of all common probes shared between EPIC v1 and EPIC v2 arrays. C) Plots showing the averaged DNA methylation beta values of common probes in the Infinium MethylationEPIC v1 array among tKD and tNN FSHD2 iPSC$^{tet\text{-}MyoD}$ samples from supplementary Fig. S3 (n= 2 and 3, respectively) and No vector samples (FSHD2 iPSC$^{tet\text{-}MyoD}$ cells without vector transfection, n= 2). Probes showing significantly increased methylation ($\Delta\beta > 0.2$ and adj.P.Val < 0.01; red) or significantly decreased methylation ($\Delta\beta < -0.2$ and adj.P.Val < 0.01; blue) are highlighted, whereas the remaining probes are shown in gray (ns). Dotted lines indicate the $\Delta\beta$ cutoffs of +0.2 and -0.2. Note that the distribution of methylation changes was relatively balanced between No vector and tNN, whereas a biased tendency toward increased methylation was observed in tKD, suggesting substantial potential off-target DNA methylation effects associated with constitutive expression of KRAB and DNMT3A-DNMT3L. D) Gene expression anlaylsis of DUX4-fl, off-target candidate genes (*SLC34A2* and *CCNA1,* identified in the volcano plot comparing tEP$^3$ vs. Neg in Fig. 7F), the myogenic marker *MYH3*, and the iPSCs marker *NANOG* in iPSCs and differentiated myocytes. Note that *SLC34A2* and *CCNA1* were expressed in differentiated FSHD2 (F2) myocyte, but were suppressed in isogenic control (IC) and non-FSHD healthy control (HC) cells. All RT-qPCR data are represented as mean±SEM. *P < 0.05, **P < 0.01, ***P < 0.001, ****P < 0.0001, and ns indicates not significant, as determined using two-way ANOVA followed by the Tukey's multiple comparisons test. E) Schematic representation of the in silico off-target prediction workflow, illustrated using a virtual sgRNA designed to target near the *MYOG* promoter as an example. The target sequence (20-mer + original PAM (AGG)) was submitted to the online CRISPRdirect tool to identify potential off-target sites with sequence matches of 20-mer, 12-mer and 8-mer with an NGG PAM, and to obtain their genomic coordinates. For tKD vs. tNN, genomic coordinates were obtained using the hg19 genome assembly because the EPICv1 array annotation is based on hg19. For LP$^3$ vs. NT, genomic coordinates were obtained using the hg38 genome assembly because EPIC v2 array annotation is based on hg38. Probes located within 500bp of each predicted off-target site were then selected, and candidate sites without nearby probes were excluded. The selected probes were subsequently mapped to annotated genes based on the gene annotation information in the corresponding array manifest file. These genes were considered potential off-target-associated genes for further analysis. Note that *MYOG* was identified in all three off-target prediction analyses based on 8-mer, 12-mer and 20-mer sequence matches. F) Numbers of potential off-target sites predicted by CRISPRdirect for sgRNAs PJ#3, PJ#6, PJ#7 and PJ#8. G) Numbers of EPIC array probes located within 500 bp of the predicted off-target sites. H) Numbers of annotated genes identified from the selected probes, representing the potential off-target-associated genes in the prediction workflow for each sgRNA.

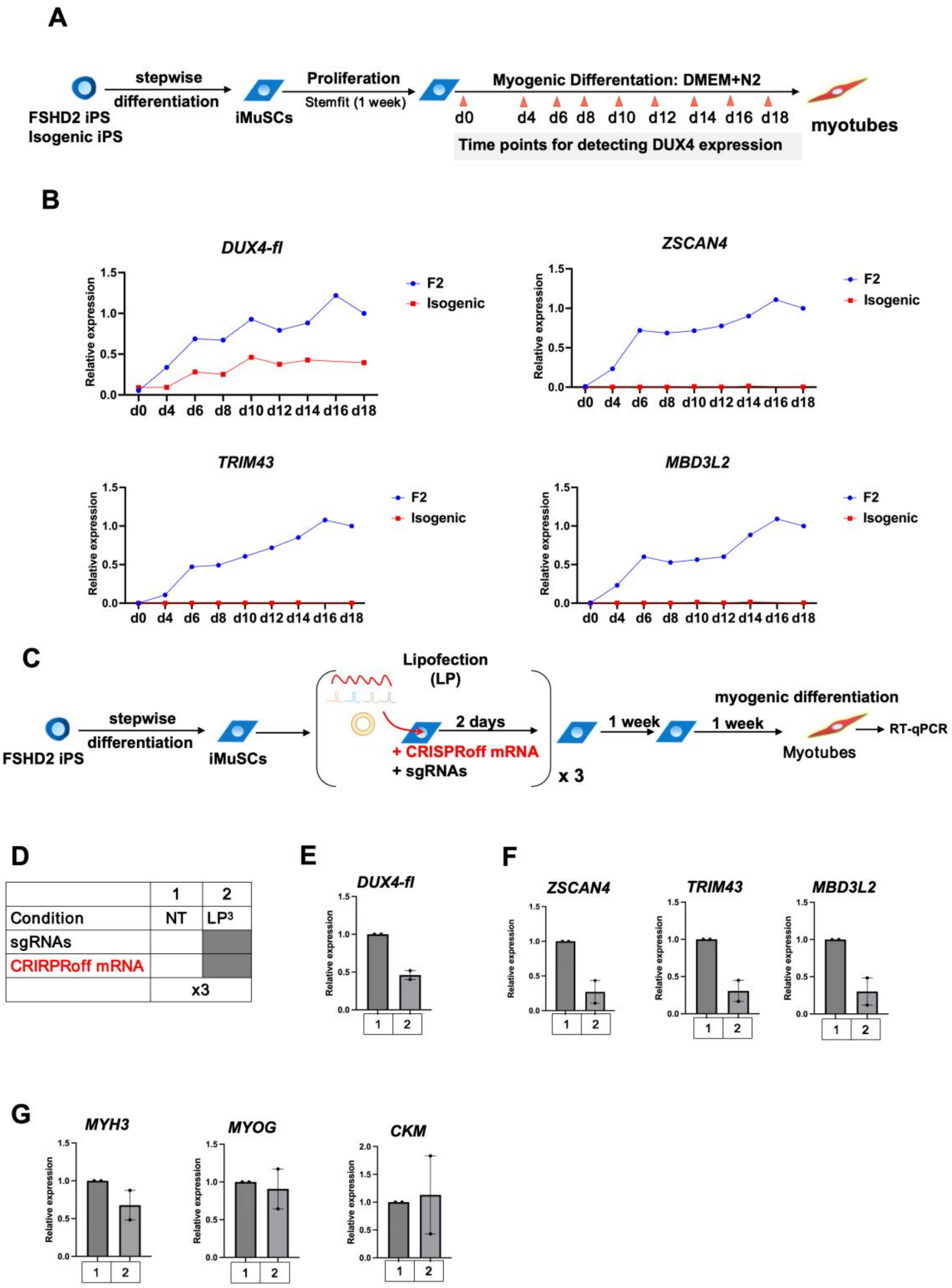


Supplementary Fig. S8. CRISPRoff-mediated hit-and-run strategy successfully suppresses *DUX4* in the FSHD2 iMuSC model.

A) Schematic representation of the generation, expansion, and myogenic differentiation of iMuSCs derived from an FSHD2 iPSC clone and its isogenic control iPSC clone. iMuSCs were generated from iPSCs through a stepwise differentiation protocol involving dermomyotome lineage induction, myogenic induction, and maturation. Following cell sorting based on the MuSC surface marker CD82, FSHD2 and isogenic control iMuSCs were isolated. After one week of expansion, myogenic differentiation was initiated. The expression levels of *DUX4* and its downstream genes were measured on day 0 and every two days from day 4 to day 18. B) Expression dynamics of *DUX4-fl* and its downstream genes (*ZSCAN4*, *TRIM43*, *MBD3L2*) during iMuSCs differentiation. Relative expression levels were normalized to those in FSHD2 iMuSCs on day18 of differentiation (n=1). C) Schematic representation of the generation, lipofection and myogenic differentiation workflow for FSHD iMuSCs. FSHD2 iMuSCs were generated from FSHD2 iPSCs through a stepwise differentiation protocol and subsequently subjected to three rounds of lipofection with CRISPRoff mRNA and sgRNAs. After one week of cell expansion, the iMuSCs were induced to differentiate into myocytes for subsequent qPCR analysis of *DUX4* and its downstream genes. D) Experimental design for comparison. FSHD2 iMuSCs were subjected to three rounds of transfection with CRISPRoff mRNA and sgRNAs ($LP^3$, conditions 2). Untreated cells (NT, condition 1) served as the negative control. E-G) qPCR analysis of *DUX4-fl* and its downstream genes (*ZSCAN4*, *TRIM43*, and *MBD3L2*), and myogenic differentiation markers (*MYH3*, *MYOG*, *CKM*), in cells on day 7 of myogenic differentiation. Relative expression were normalized to those of the untreated control (NT, condition1) within each experimental set (n=2).
